# Barnacle-Inspired Paste for Instant Hemostatic Tissue Sealing

**DOI:** 10.1101/2020.12.12.422505

**Authors:** Hyunwoo Yuk, Jingjing Wu, Xinyu Mao, Claudia E. Varela, Ellen T. Roche, Christoph S. Nabzdyk, Xuanhe Zhao

## Abstract

Whilst sealing damaged tissues by adhesives has potential advantages over suturing or stapling, existing tissue adhesives cannot form rapid or robust adhesion on tissues covered with body fluids such as blood. In contrast, the glues of barnacles, consisting of a lipid-rich matrix and adhesive proteins, and can strongly adhere to wet and contaminated surfaces. Here we report a barnacle-inspired paste capable of forming instant robust hemostatic sealing of diverse tissues. The paste is composed of a hydrophobic oil matrix and bioadhesive microparticles to implement the barnacle-inspired mechanism to repel blood through the hydrophobic matrix. Subsequently, the bioadhesive microparticles crosslink with underlying tissues under gentle pressure. The barnacle-inspired paste can provide tough (interfacial toughness over 300 J m^-2^) and strong (shear and tensile strength over 70 kPa, burst pressure over 350 mmHg) hemostatic sealing of a broad range of tissues within five seconds. We validate *in vitro* and *in vivo* biocompatibility and biodegradability of the barnacle-inspired paste in rodent models. We further demonstrate potential applications of the barnacle-inspired paste for instant hemostatic sealing in *ex vivo* porcine aorta, *in vivo* rat liver and heart models.

---

Tissue and organ-related hemorrhage can be life-threatening and are challenging to treat due to their highly time-sensitive and possibly complex nature^1^. For example, uncontrolled hemorrhages is one of the major causes of mortality in the world, accounting for over two million deaths annually^2,3^. Existing topical hemostatic agents mostly aim to augment and accelerate intrinsic blood coagulation to achieve hemostasis^4–10^. This is either achieved by coagulation factor concentration by rapid water absorption or local procoagulant delivery^3,9^. However, hemostasis through blood coagulation cannot yield immediate hemorrhage control due to the inherently gradual nature of blood clot formation^9^. Furthermore, rapid or pressurized blood flow through a wound bed can wash out any forming blood clot, potentially limiting the efficacy and duration of hemostasis. Furthermore, coagulation-dependent hemostasis mechanisms are less effective in anticoagulated or coagulopathic patients^11^.

In contrast, adhesive sealing of bleeding tissues offers a promising alternative to blood coagulation for hemostasis^4–8^, but existing tissue adhesives display several substantial limitations. Commercially-available tissue adhesives provide only weak and/or slow adhesion formation with tissue surfaces covered by blood^9,12,13^. Whilst a few blood-resistant tissue adhesives with improved adhesion performance have been developed, the need for ultraviolet (UV) irradiation^12–14^ and/or prolonged steady pressure application (e.g., over 3 min)^15^ to form adhesion substantially limits their utility for clinical applications.

In nature, marine invertebrates such as barnacles^16–19^, mussels^20–22^, and sandcastle worms^23^ show a remarkable capability of forming strong adhesion on wet and contaminated surfaces. In particular, barnacles can strongly adhere to a broad range of underwater surfaces, ranging from man-made structures to animal skins (**Fig. 1a**)^17,24^. Glues of barnacles are known to consist of two major components: a lipid-rich matrix and adhesive proteins which synergistically offer strong adhesion on wet and contaminated surfaces (**Fig. 1b,c**)^16,24,25^. The lipid-rich matrix of the barnacle glues first cleans the underlying substrate by repelling water and contaminants (**Fig. 1b**), and subsequently the adhesive proteins crosslink with the substrate to form stable and strong adhesion (**Fig. 1c**)^24,25^. Whilst the adhesive proteins of the barnacle glues have been investigated and synthesized to develop underwater and tissue adhesives^21,26,27^, the aforementioned mechanism – based on the synergistic interplay between the lipid-rich matrix and adhesive proteins in the barnacle glues – has remained unexplored for tissue adhesives.

**Fig. 1.**
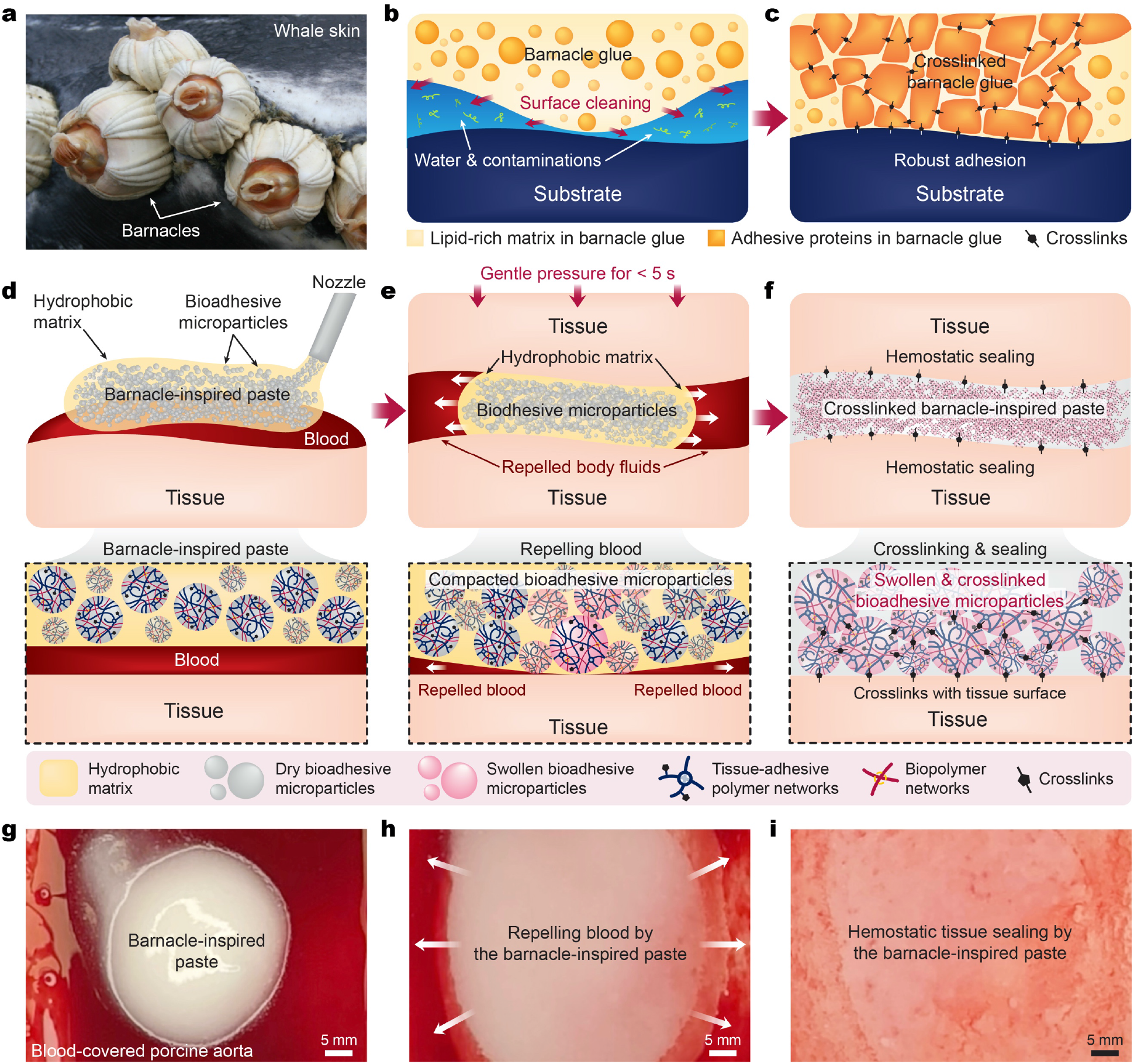
Design and mechanism of the barnacle-inspired paste. **a**, Barnacles adhered to the skin of a whale. The photo is credited to Aleria Jensen, the U.S. National Oceanic and Atmospheric Administration (NOAA). **b,c**, Composition and adhesion mechanism of the barnacle glue on substrates covered with water and contaminants. A lipid-rich matrix in the barnacle glue cleans the substrate by repelling water and contaminants (b) while the adhesive proteins can crosslink and form robust adhesion on the cleaned substrate (c). **d-f**, Design of a barnacle-inspired paste consisting of bioadhesive microparticles and a hydrophobic oil matrix (d). Repel-crosslinking mechanism of the barnacle-inspired paste integrates repelling of blood (e) and subsequent formation of hemostatic sealing by crosslinking (f). **g-i**, Photographs of the barnacle-inspired paste injected on blood-covered porcine aorta (g), pressed with a gelatin-coated glass substrate (i), and formed hemostatic tissue sealing (i), corresponding to each panel in (d-f).

Here we propose a barnacle-inspired bioadhesive to achieve instant robust hemostatic sealing of diverse tissues. The barnacle-inspired bioadhesive takes the form of an injectable paste consisting of a hydrophobic oil matrix and bioadhesive microparticles (**Fig. 1d,g**), which take on similar functional roles as the lipid-rich matrix and the adhesive proteins in barnacle glues, respectively. Upon application of gentle pressure (e.g., 10 kPa), the hydrophobic oil matrix repels the blood allowing the compacted bioadhesive microparticles to interact with one another and with the tissue surface underneath (**Fig. 1e,h**). Subsequently, the compacted bioadhesive microparticles crosslink with one other and with the tissue surface to form robust adhesion within 5 s without the need for additional adjuncts such as UV irradiation or coagulation of blood (**Fig. 1f,i**, **Supplementary Figs. 1,2**, and **Supplementary Video 1**).

## Results and Discussion

### Design and mechanism of the barnacle-inspired paste

To implement the proposed design of the barnacle-inspired paste, we first prepare the bioadhesive microparticles by cryogenically grinding a dry bioadhesive sheet (**Supplementary Fig. 3**)^28^. The resultant bioadhesive microparticles are composed of crosslinked networks of poly(acrylic acid) grafted with *N*-hydroxysuccinimide ester (PAA-NHS ester) and chitosan, which provide bioadhesiveness and robust mechanical properties, respectively (**Supplementary Fig. 4**)^28^. The barnacle-inspired paste is then prepared by thoroughly mixing the bioadhesive microparticles with a biocompatible hydrophobic matrix such as silicone oil and essential oil to form a stable paste (See **Supplementary Text** for a detailed discussion on stability of the barnacle-inspired paste, **Supplementary Fig. 3**)^29–31^.

Notably, the average size of the bioadhesive microparticles can be controlled by the cryogenic grinding conditions. When keeping the grinding time constant at 2 minutes and changing the grinding frequency ranging from 10 Hz to 30 Hz, we find that a higher grinding frequency results in a smaller average size of bioadhesive microparticles (∼ 200 µm at 10 Hz and ∼ 10 µm at 30 Hz, **Supplementary Fig. 5**). Since the smaller bioadhesive microparticles result in better injectability of the barnacle-inspired paste and its ability to conform complex geometry of bleeding tissues (**Supplementary Fig. 6**), we choose the grinding frequency of 30 Hz in this study. Furthermore, the rheological property of the barnacle-inspired paste can be tuned by controlling the ratio between the bioadhesive microparticles and the hydrophobic oil matrix^32^. By increasing the volume fraction *ϕ* of the bioadhesive microparticles, the barnacle-inspired paste exhibits a transition from a viscous fluid (*ϕ* < 0.3) to a stable thixotropic paste (*ϕ* > 0.4) (See **Supplementary Text** for a detailed discussion on rheological property of the barnacle-inspired paste, **Supplementary Fig. 7**). An injectable thixotropic paste is more favorable than an easily flowing viscous liquid to provide more reproducible and stable application of the barnacle-inspired paste to the bleeding target injury. Hence, we choose a relatively high-volume fraction of the bioadhesive microparticles (*ϕ* = 0.4) in this study.

To investigate the influence of the hydrophobic oil matrix and the bioadhesive microparticles on hemostatic sealing, we measure the pull-off force between porcine heart sealed by the bioadhesive microparticles with various matrix materials (**Fig. 2a**). The pull-off tests performed both in air and a Dulbecco’s Modified Eagle Medium (DMEM) bath show similarly high pull-off forces (*p* = 0.25 for air; *p* = 0.49 for DMEM) between the bioadhesive microparticles without and with silicone oil matrix. In contrast, silicone oil without the bioadhesive microparticles and the bioadhesive microparticles with hydrophilic polyethylene glycol (PEG) matrix exhibit significantly lower pull-off force (**Fig. 2b,c**). These results indicate that the bioadhesive microparticles alone can provide instant robust sealing between wet tissues in the absence of blood.

**Fig. 2.**
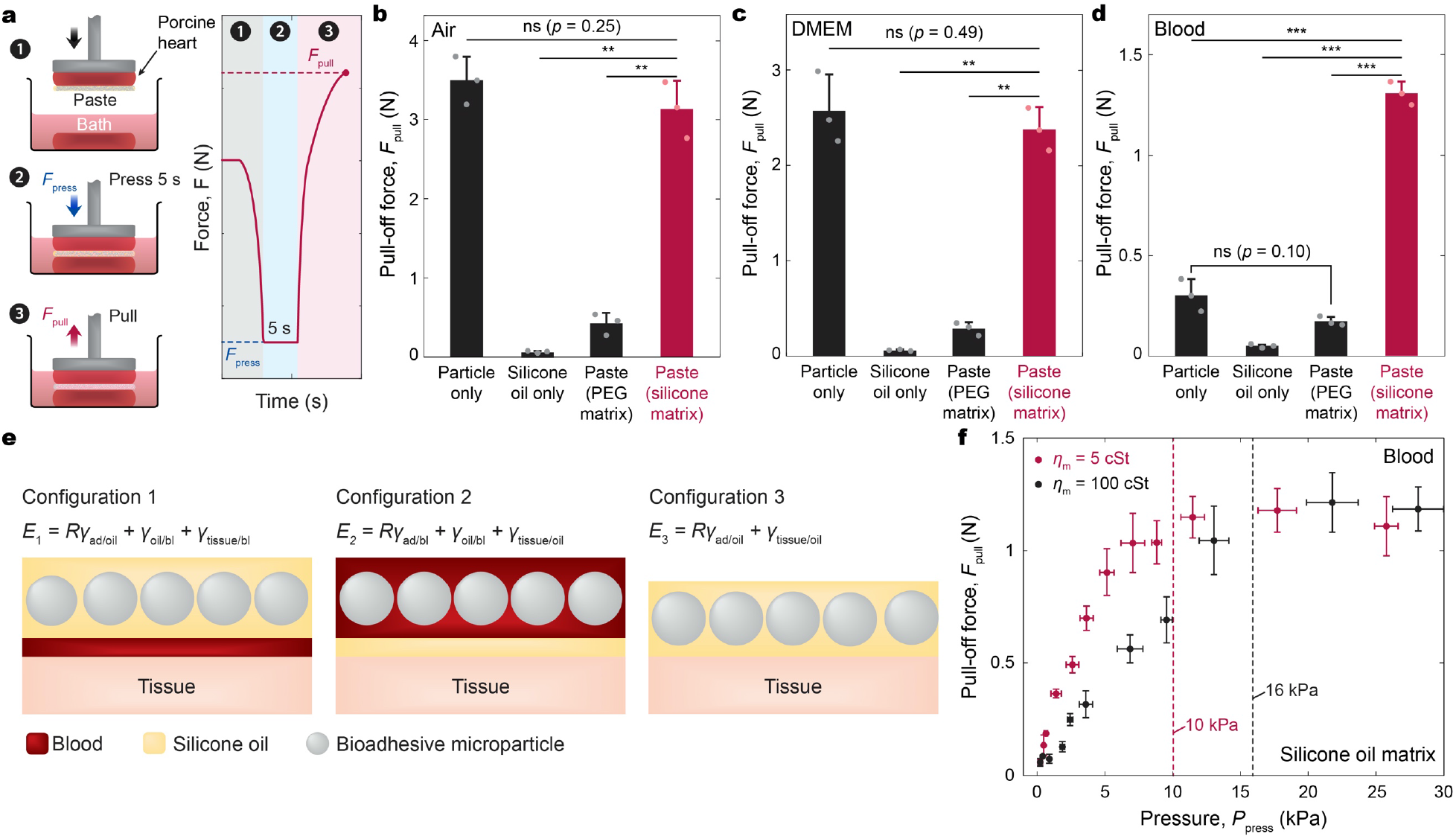
Effects of matrix materials in the barnacle-inspired paste. **a**, Schematic illustrations of the setup and procedure for pull-off tests. The surface area of porcine heart is 1 cm^2^. *F*_press_, pressing force; *F*_pull_, pull-off force. **b-d**, Pull-off forces between tissues sealed by the bioadhesive microparticles, silicone oil, the bioadhesive microparticles with PEG matrix, and the bioadhesive microparticles with silicone oil matrix measured in air (b), DMEM (c), and heparinized porcine blood bath (d). DMEM, Dulbecco’s Modified Eagle Medium. **e**, Schematic illustrations of configurations for the barnacle-inspired paste and the corresponding total surface energies. *E*, the total surface energy of each configuration; *R*, the roughness factor representing the ratio of the actual and projected surface areas of the bioadhesive microparticles; *γ*_A/B_, the interfacial energy between A and B (subscript “ad” represents the bioadhesive microparticle and “bl” represents blood). **f**, Pull-off forces vs. applied pressure between tissues sealed by the bioadhesive microparticles with varying viscosity silicone oil matrices (5 cSt or 100 cSt) measured in heparinized porcine blood bath. Vertical dashed lines indicate threshold applied pressures. *η*_m_, kinematic viscosity of the silicone oil. Values in **b**-**d,f** represent the mean and the standard deviation (*n* = 3). Statistical significance and *p* values are determined by two-sided Student *t*-test; ns, not significant; ** *p* ≤ 0.01; *** *p* ≤ 0.001.

It is also notable that hydrophilic matrix materials such as PEG can significantly compromise the adhesive performance of the bioadhesive microparticles by prematurely swelling the bioadhesive microparticles before contacting the wet tissue surface^28,33^. The pull-off tests performed in a heparinized porcine blood bath display a high pull-off force for the bioadhesive microparticles with the silicone oil matrix, whilst the bioadhesive microparticles without the silicone oil matrix exhibit significantly the lower pull-off forces (**Fig. 2d**). In the absence of a protective hydrophobic matrix, the blood can readily infiltrate into and interact with the bioadhesive microparticles, preventing the formation of robust adhesion among the microparticles or with the tissue surface (**Supplementary Fig. 8**). These results indicate that the silicone oil matrix can effectively protect and preserve the bioadhesive microparticles and their adhesive capabilities in the presence of blood.

To further understand the silicone oil’s role as a protective matrix for the bioadhesive microparticles against blood, we compare the total surface energies of three configurations: i) the bioadhesive microparticles completely wetted by the silicone oil with a layer of blood facing below it (protected state, *E*_1_), ii) the bioadhesive microparticles completely wetted by blood with a layer of the silicone oil facing below it (unprotected state, *E*_2_), and iii) the bioadhesive microparticles completely wetted by the silicone oil without blood (repelled state, *E*_3_)^34,35^ (**Fig. 2e**). To ensure the energetically stable protection of the bioadhesive microparticles and the repelling of blood by the silicone oil matrix, one should satisfy Δ*E*_A_ = *E*_2_ − *E*_1_ > 0 and Δ*E*_B_ = *E*_1_ − *E*_3_ > 0, respectively, which can be expressed as

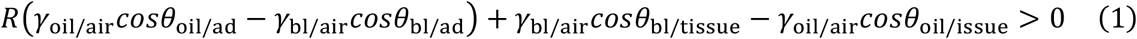

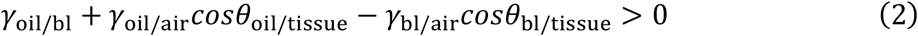

where *R* is the roughness factor representing the ratio of the actual and projected surface areas of the bioadhesive microparticles, *γ*_A/B_ represents the interfacial energy between A and B, and *θ*_A/B_ represents the contact angle of A on B (subscript “ad” represents the bioadhesive microparticle; “bl” represents blood). Note that we take *R* approximately as *π* for the bioadhesive microparticles in the barnacle-inspired paste based on the approximation of tightly placed spherical particles with the same diameter. By substituting the values of parameters (*R* = *π, γ*_oil/air_ = 20.9 mN m^−1^, *γ*_bl/air_ = 72.0 mN m^−1^, *γ*_oil/bl_ = 40 mN m^−1^, *θ*_oil/ad_ = 4.5°, *θ*_bl/ad_ = 96°, *θ*_oil/tissue_ = 4.2°, *θ*_bl/tissue_ = 84°, see **Methods** for details) into Eqs. (1–2), it is evident that the barnacle-inspired paste satisfies Eqs. (1) and (2). Hence, the silicone oil matrix can protect the bioadhesive microparticles against blood and repel blood from the tissue surface, supporting our proposed design and mechanism (**Fig. 1d-f** and **Supplementary Fig. 2**).

### Adhesion mechanism and performance

Next, we further investigate the behavior of bioadhesive microparticles in the process of adhering to the tissue surface. Upon application of gentle pressure, the bioadhesive microparticles in the barnacle-inspired paste are densely compacted and form a non-flowable jammed adhesive layer (**Supplementary Fig. 9**). During this compaction process, the silicone oil matrix is pushed out of the barnacle-inspired paste, repelling and cleaning blood from the tissue surface (**Supplementary Fig. 10** and **Supplementary Video 2**). Hence, the applied pressure and the property of the silicone oil matrix can affect the adhesion formation of the barnacle-inspired paste. To investigate the effects of the applied pressure and the property of the silicone oil matrix on adhesion to tissues, we measure the pull-off force between porcine heart sealed by the barnacle-inspired paste with varying applied pressure as well as viscosity of the silicone oil matrix (**Fig. 2f**). As the applied pressure increases, the pull-off force first increases and then reaches a plateau above a certain threshold of applied pressure (e.g., 10 kPa for silicone oil with 5 cSt kinematic viscosity) (**Fig. 2f**). Moreover, the silicone oil with a higher viscosity shows a higher threshold of the applied pressure (e.g., 16 kPa for silicone oil with 100 cSt kinematic viscosity) (**Fig. 2f**), agreeing with the granular suspension theory in the literature^36^ (See **Supplementary Text** for a detailed discussion on compaction of the barnacle-inspired paste).

Once the barnacle-inspired paste forms a densely compacted layer on the wet tissue surface, the bioadhesive microparticles in the barnacle-inspired paste establish instant physical crosslinks between themselves and to the tissue surface by hydrogen bonds between carboxylic acid and primary amine groups abundant in the bioadhesive microparticles and biological tissues (**Supplementary Figs. 2** and **11**). The physically crosslinked bioadhesive microparticles further swell in wet physiological environment (**Supplementary Fig. 12**) and subsequently form stable covalent crosslinks between NHS ester groups in the bioadhesive microparticles and primary amine groups abundant in both biological tissues and chitosan in the bioadhesive microparticles within a few min^37^ (**Supplementary Figs. 2** and **13**). Notably, the bioadhesive microparticles rapidly swell (**Supplementary Fig. 12a,b**) and become soft hydrogels whose Young’s modulus is similar to that of soft tissues (**Supplementary Fig. 14**) in contact with wet tissue surfaces. This rapid transition to soft hydrogels in contact with the wet tissue surfaces may minimize potential tissue damages during application process of the barnacle-inspired paste.

To evaluate the adhesion performance of the barnacle-inspired paste, we conduct three different types of mechanical tests to measure interfacial toughness (180-peel tests, ASTM F2256), shear strength (lap-shear tests, ASTM F2255), and tensile strength (tensile tests, ASTM F2258), respectively (**Supplementary Fig. 15**). The barnacle-inspired paste forms robust adhesion (interfacial toughness over 240 J m^-2^) between blood-covered porcine skin after gentle pressure (10 kPa) application for 5 s, demonstrating the instant blood-resistant sealing capability (**Fig. 3a**). The tissues sealed created by the barnacle-inspired paste shows no significant deterioration in the measured interfacial toughness (*p* = 0.78) over 48 h after the initial pressing (**Fig. 3b**). The barnacle-inspired pate demonstrates superior adhesion performance compared to existing commercially-available tissue adhesives, including gelatin-based hemostatic adhesives (Surgiflo®), fibrin-based hemostatic adhesives (Tisseel, TachoSil®), albumin-based adhesives (BioGlue®), PEG-based adhesives (Coseal), and cyanoacrylate adhesives (Histoacryl®). These commercially-available tissue adhesives exhibit relatively slow adhesion formation (longer than 1 min) and limited adhesion performance on blood-covered porcine skin (interfacial toughness lower than 30 J m^-2^ and shear/tensile strength lower than 20 kPa) (**Fig. 3c-e**). In contrast, the barnacle-inspired paste can form instant robust adhesion with interfacial toughness over 240 J m^-2^, shear strength over 70 kPa, and tensile strength over 50 kPa within 5 s, substantially outperforming the commercially-available tissue adhesives and glues (**Fig. 3c-e**). Furthermore, the barnacle-inspired paste is applicable for a broad range of tissues covered by blood to form robust adhesion with high interfacial toughness (over 240 J m^-2^ for skin, 150 J m^-2^ for aorta, 140 J m^-2^ for heart, 330 J m^-2^ for muscle), shear and tensile strength (over 70 kPa for skin, 55 kPa for aorta, 45 kPa for heart, 50 kPa for muscle) in less than 5 s (**Fig. 3f-h** and **Supplementary Fig. 16**). Furthermore, the barnacle-inspired paste can form instant robust sealing of tissues covered by other body fluids such as mucus and gastric juice, potentially offering a broader applicability for sealing of various tissues beyond the clinical context of hemostasis (**Fig. 3f-h** and **Supplementary Fig. 16**). Notably, the barnacle-inspired glue’s superior adhesion performance can also be achieved based on various hydrophobic matrix materials other than silicone oil such as soybean oil (**Supplementary Fig. 17**).

**Fig. 3.**
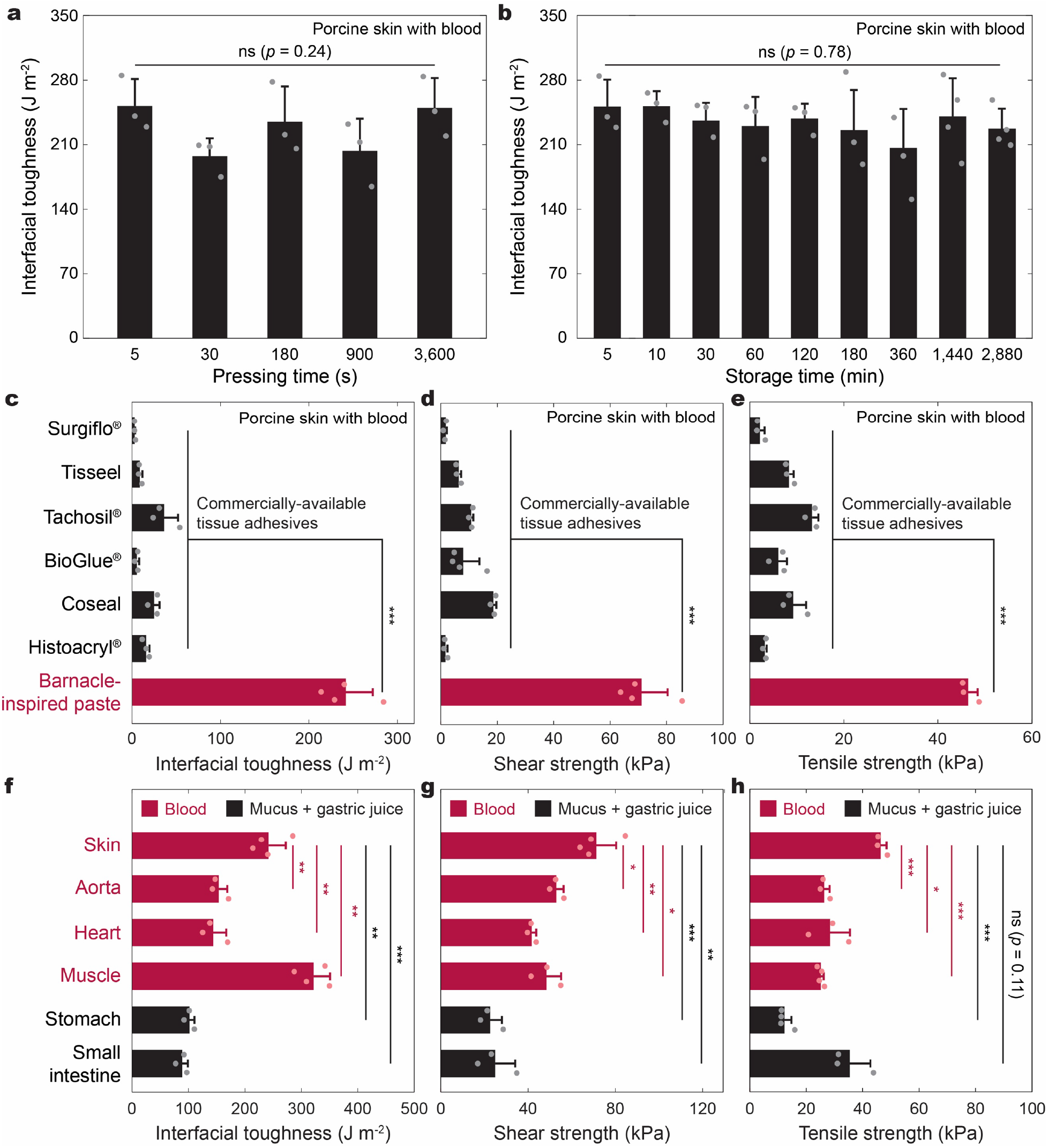
Adhesion performance of the barnacle-inspired paste. **a**, Interfacial toughness vs. pressing time of blood-covered porcine skin sealed by the barnacle-inspired paste. **b**, Interfacial toughness vs. storage time of blood-covered porcine skin sealed by the barnacle-inspired paste. **c-e**, Comparison of interfacial toughness (c), shear strength (d), and tensile strength (e) of blood-covered porcine skin sealed by the barnacle-inspired paste and various commercially-available products. **f-h**, Interfacial toughness (f), shear strength (g), and tensile strength (h) of various tissues covered by blood or mucus sealed by the barnacle-inspired paste. Values represent the mean and the standard deviation (*n* = 3-4). Statistical significance and *p* values are determined by one-way ANOVA and Tukey’s multiple comparison test; ns, not significant; * *p* ≤ 0.05; ** *p* ≤ 0.01; *** *p* ≤ 0.001.

### Biocompatibility and biodegradability

To evaluate biocompatibility and biodegradability of the barnacle-inspired paste, we perform *in vitro* and *in vivo* characterizations based on rat models (**Fig. 4**). A cell culture media (DMEM) conditioned with the barnacle-inspired paste exhibits comparable *in vitro* cytotoxicity of rat cardiomyocytes to a control (pristine DMEM) after 24-hour culture (**Fig. 4a**). We evaluate *in vivo* biocompatibility of the barnacle-inspired paste based on dorsal subcutaneous implantation in rat models for various time points from 1 day to 2 weeks (**Fig. 4b,c** and **Supplementary Fig. 18a,b**). Histological assessment by a blinded pathologist demonstrates that the barnacle-inspired paste elicits very mild to mild inflammation comparable to a commercially-available U.S. Food and Drug Administration (FDA)-approved tissue adhesive Coseal at all time points (*p* = 0.39 for 1 day, *p* = 0.54 for 3 days, *p* = 0.67 for 1 week, *p* = 0.21 for 2 weeks, **Fig. 4d**). To further investigate *in vivo* biocompatibility of the barnacle-inspired paste, we perform immunofluorescence staining of various markers for fibroblast (αSMA), type 1 collagen (Collagen I), T-cell (CD3), and macrophage (CD68) related to inflammatory and foreign body responses (**Fig. 4e-h** and **Supplementary Fig. 18c-f**). The normalized immunofluorescence intensity analysis shows that the barnacle-inspired paste induces no significant difference in the expression of αSMA, Collagen I, CD3, and CD68 compared to that of Coseal for all time points (**Fig. 4i-l**). Notably, the barnacle-inspired paste exhibits gradual biodegradation via resorption by macrophages over a longer period of implantation (**Supplementary Fig. 19a-c**), and the rate of biodegradation can be accelerated by using faster degrading materials such as gelatin to replace chitosan in the bioadhesive microparticles (**Supplementary Fig. 19d**).

**Fig. 4.**
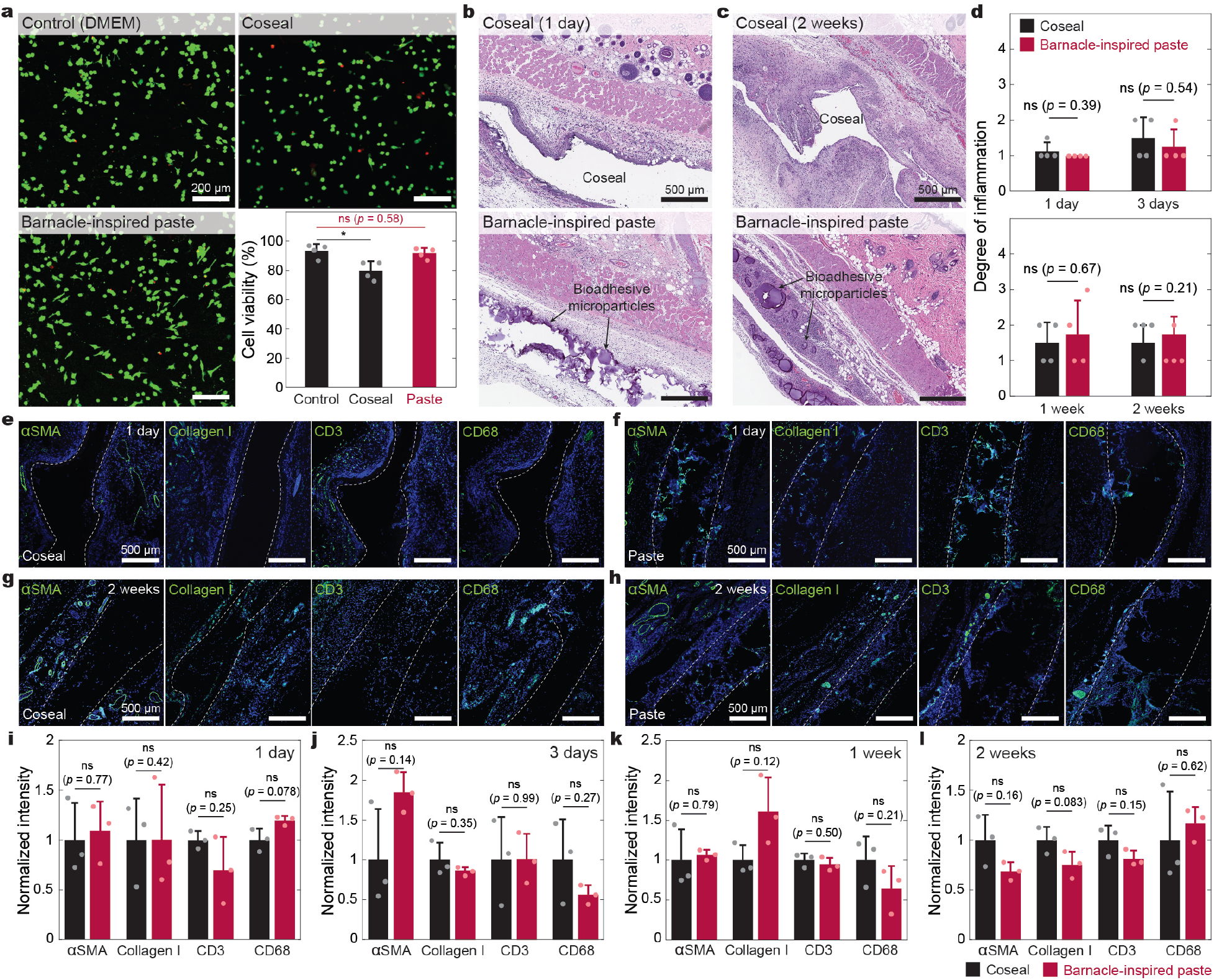
Biocompatibility of the barnacle-inspired paste. **a**, *In vitro* cell viability of rat cardiomyocytes based on LIVE/DEAD assay after 24-h culture in control media (DMEM), Coseal-incubated media, and the barnacle-inspired paste-incubated media (bottom-right panel). Representative confocal images of the LIVE/DEAD assay for control (top-left panel), Coseal (top-right panel), and the barnacle-inspired paste (bottom-left panel). DMEM, Dulbecco’s Modified Eagle Medium. **b,c**, Representative histology images stained with hematoxylin and eosin (H&E) for Coseal and the barnacle-inspired paste after rat subcutaneous implantation for 1 day (b) and 2 weeks (c). 4 independent experiments were conducted with similar results. **d**, Degree of inflammation evaluated by a blinded pathologist (0, normal; 1, very mild; 2, mild; 3, severe; 4, very severe). **e-h**, Representative immunofluorescence images of Coseal (e) and the barnacle-inspired paste (f) after rat subcutaneous implantation for 1 day; Coseal (g) and the barnacle-inspired paste (h) after rat subcutaneous implantation for 2 weeks. Cell nuclei are stained with 4′,6-diamidino-2-phenylindole (DAPI, Blue). Green fluorescence corresponds to the expression of fibroblast (αSMA), type 1 collagen (Collagen-I), T-cell (CD3), and macrophage (CD68), respectively. **i-l**, Normalized fluorescence intensity from the immunofluorescence images for Coseal and the barnacle-inspired paste after rat subcutaneous implantation for 1 day (i), 3 days (j), 1 week (k), and 2 weeks (l). Values in **a,d,i-l** represent the mean and the standard deviation (*n* = 4 independent samples). Statistical significance and *p* values are determined by two-sided Student *t*-test; ns, not significant; * *p* ≤ 0.05.

### Instant hemostatic tissue sealing

The barnacle-inspired paste’s unique capability of forming instant robust adhesion on blood-covered tissues can be advantageous for rapid and coagulation-independent hemostatic sealing of various tissues in clinical and biomedical applications. To evaluate instant hemostatic tissue sealing capability of the barnacle-inspired paste, we demonstrate hemostatic sealing in models of *ex vivo* porcine aortic and *in vivo* rat liver and heart injuries. Within 5s, the barnacle-inspired paste can form instant hemostatic sealing of a bleeding *ex vivo* porcine aorta (3-mm hole; blood flow pressure at 150 mmHg) through which an anticoagulated (heparinized) porcine blood bath is circulated (**Supplementary Fig. 20a** and **Supplementary Video 3**). The aorta, sealed by the barnacle-inspired paste, maintains a seal under continuous blood flow and can withstand a supraphysiological pressure (250 mmHg) without leakage 6 h after the application and under continuous flow (**Supplementary Video 3**). Importantly, filtering of the blood bath with a mesh (100-µm mesh size) after the test (closed-loop flow for 6 h) does not yield any evidence of released free-floating bioadhesive microparticles. This demonstrates the robust hemostatic sealing by the barnacle-inspired paste and a low potential risk of microparticle embolization (**Supplementary Fig. 20b**). We further quantitatively evaluate the strength of hemostatic sealing based on a standard burst pressure measurement (ASTM F2392-04). The barnacle-inspired paste shows a burst pressure over 350 mmHg, significantly exceeding the normal systolic arterial blood pressure in human (∼ 120 mmHg) and the burst pressure of hemostatic sealing made by sutures and commercially-available hemostatic sealants (Surgiflo®, Tisseel, Tachosil®) (**Supplementary Fig. 20c**).

To quantitatively evaluate the hemostatic sealing capability of the barnacle-inspired paste *in vivo*, we measure the time to hemostasis and blood loss until hemostasis based on rat hepatic and cardiac hemostasis models (**Figs. 5** and **6**). The barnacle-inspired paste can form hemostatic sealing of bleeding liver (5-mm diameter and 2-mm depth injury) and heart (2-mm diameter ventricular wall injury) within 5 s (**Figs. 5a** and **6a**), giving significantly less time to hemostasis (**Figs. 5c** and **6c**) and blood loss until hemostasis (**Figs. 5d** and **6d**) compared to the injury group (without hemostasis) and commercially-available hemostatic agents Surgicel and Coseal (**Supplementary Figs. 21** and **22**). Notably, the injury, Surgicel, and Coseal groups are not able to form hemostatic sealing of heart due to the high pressure and high volume bleeding from a ventricular injury (**Fig. 6c,d, Supplementary Fig. 22**, and **Supplementary Video 4**), whereas the barnacle-inspired paste can form instant hemostatic sealing of a ventricular injury within 5 s, restoring the normal intraventricular blood pressure immediately after hemostasis (**Fig. 6e** and **Supplementary Video 5**).

We also demonstrate that the barnacle-inspired paste maintains a seal on the injured liver and heart 2 weeks after the initial hemostatic application (**Figs. 5b** and **6b**). Furthermore, histological analysis of the sealed liver and heart tissues indicates that the barnacle-inspired paste allows cell infiltration into the crosslinked bioadhesive microparticles and healing of the underlying injuries (**Figs. 5e** and **6f,g**). Immunofluorescence analysis of inflammatory and foreign body response markers (CD3 for T-cell; CD68 for macrophage) shows that the barnacle-inspired paste induces the expression of CD3 and CD68 comparable to that of Coseal and significantly lower than that of Surgicel (**Fig. 5f-i**). Furthermore, complete blood count (CBC) and blood chemistry analysis of the animals 2 weeks after hemostatic sealing demonstrates that the barnacle-inspired paste does not show significant difference in inflammation-related blood cells and markers for organ-specific diseases compared to the healthy animals (**Supplementary Figs. 23** and **24**).

**Fig. 5.**
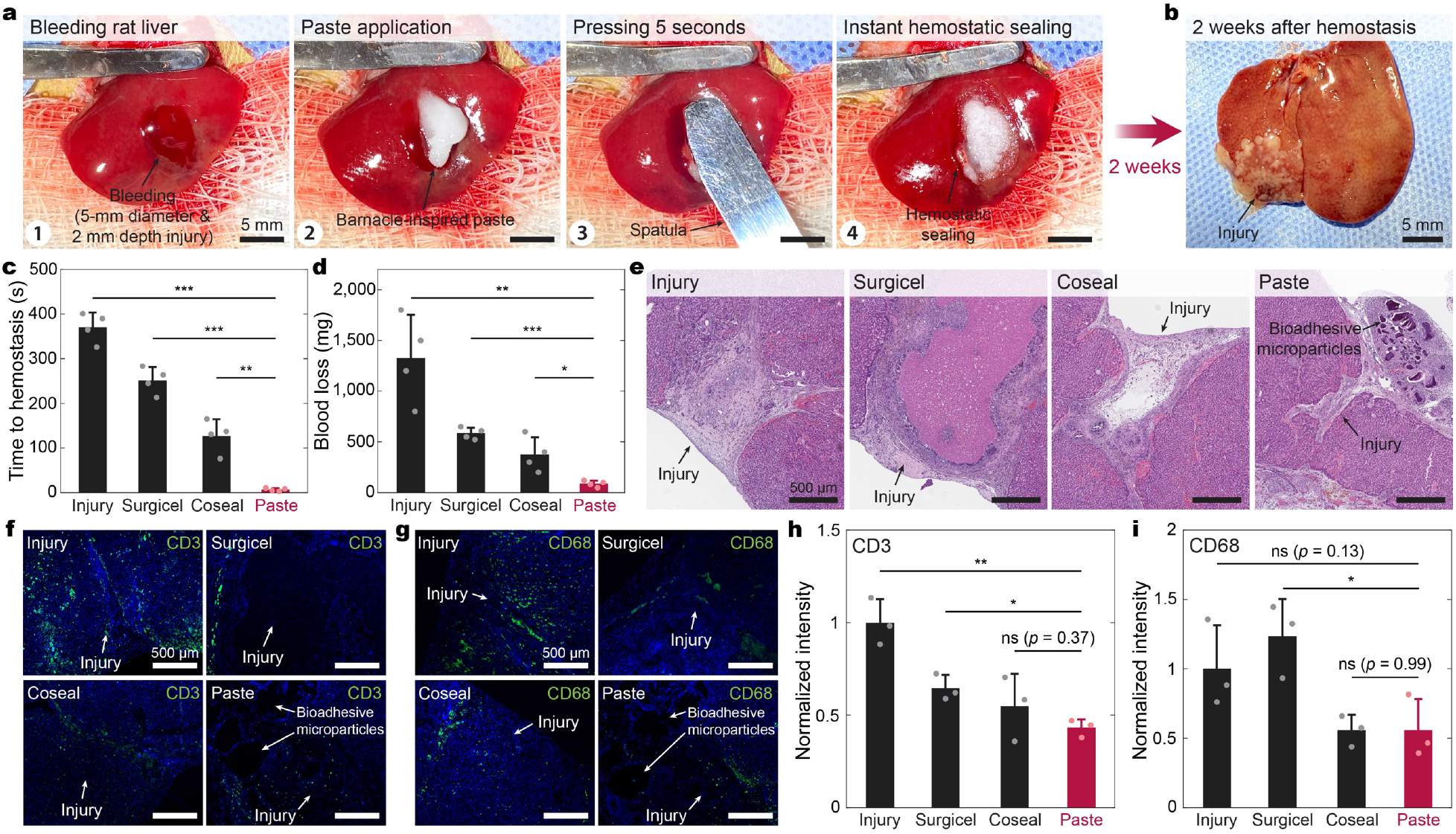
Instant hemostatic sealing of liver by the barnacle-inspired paste. **a**, Instant hemostatic sealing of a bleeding rat liver *in vivo* by the barnacle-inspired paste. **b**, Excised rat liver 2 weeks after hemostatic sealing by the barnacle-inspired paste. **c,d**, Time to hemostasis (c) and blood loss until hemostasis (d) for hepatic bleeding without treatment (injury), Surgicel, Coseal, and the barnacle-inspired paste. **e**, Representative histology images stained with hematoxylin and eosin (H&E) for injured liver without treatment (injury) and hemostatic sealing formed by Surgicel, Coseal, and the barnacle-inspired paste 2 weeks after hemostasis. 4 independent experiments were conducted with similar results. **f,g**, Representative immunofluorescence images for injured liver without treatment (injury) and with hemostatic sealing by Surgicel, Coseal, and the barnacle-inspired paste 2 weeks after hemostasis. Cell nuclei are stained with 4′,6-diamidino-2-phenylindole (DAPI, Blue). Green fluorescence corresponds to the expression of T-cell (CD3, f) and macrophage (CD68, g), respectively. **h,i**, Normalized fluorescence intensity from the immunofluorescence images for CD3 (h) and CD68 (i). Values in **c,d,h,i** represent the mean and the standard deviation (*n* = 4 independent samples). Statistical significance and *p* values are determined by two-sided Student *t*-test; ns, not significant; * *p* ≤ 0.05; ** *p* ≤ 0.01; *** *p* ≤ 0.001.

**Fig. 6.**
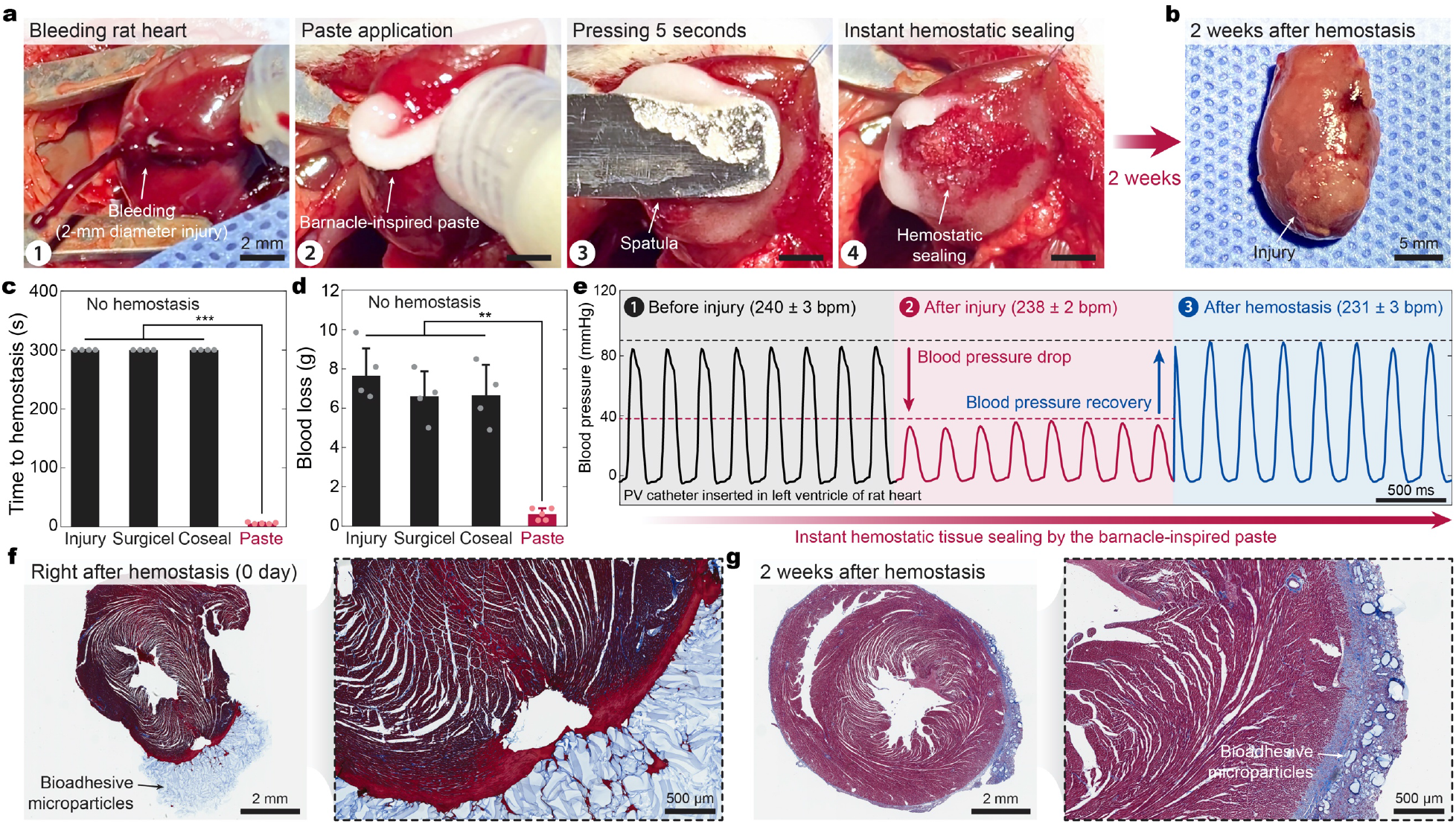
Instant hemostatic sealing of heart by the barnacle-inspired paste. **a**, Instant hemostatic sealing of a bleeding rat heart *in vivo* by the barnacle-inspired paste. **b**, Excised rat heart 2 weeks after hemostatic sealing by the barnacle-inspired paste. **c,d**, Time to hemostasis (c) and blood loss until hemostasis (d) for cardiac bleeding without treatment (injury), Surgicel, Coseal, and the barnacle-inspired paste. **e**, Intraventricular blood pressure and heart rate of a rat heart before injury, after injury (2-mm biopsy punch), and after hemostatic sealing by the barnacle-inspired paste. bpm, beats per minute. **f,g**, Representative histology images stained with Masson’s trichrome for injured heart with hemostatic sealing by the barnacle-inspired paste right after (f) and 2 weeks after (g) hemostasis. 4 independent experiments were conducted with similar results. Values in **c,d** represent the mean and the standard deviation (*n* = 4 independent samples). Statistical significance and *p* values are determined by one-way ANOVA and Tukey’s multiple comparison test; ** *p* ≤ 0.01; *** *p* ≤ 0.001.

In summary, we report a barnacle-inspired paste that incorporates a unique biologically-inspired design and adhesion mechanism to address the limitations of existing hemostatic agents and tissue adhesives. This study provides an effective tool for instant and coagulation-independent hemostasis, wound closure, and surgical repair, potentially offering a promising solution to life-threatening critical clinical challenges. The barnacle-inspired repel-crosslinking mechanism may further offer valuable insights for the design and development of adhesives in wet and contaminated environments.

## Methods

### Materials

All chemicals were obtained from Sigma-Aldrich unless otherwise mentioned and used without further purification. For preparation of dry bioadhesive, acrylic acid, acrylic acid *N*-hydroxysuccinimide ester (AAc-NHS ester), α-ketoglutaric acid, chitosan (HMC+ Chitoscience Chitosan 95/500, 95 % deacetylation), and gelatin (type A bloom 300 from porcine skin) were used. For matrix of bioadhesive microparticles, silicone oils with different viscosity (5 cSt and 100 cSt), soybean oil, or polyethylene glycol (PEG) were used. For visualization of the barnacle-inspired paste, fluorescein-labeled chitosan (KITO-8, PolySciTech®) was used for confocal microscope images. Heparinized porcine blood was purchased from Lampire Biological Laboratories, Inc. All porcine tissues and organs for *ex vivo* experiments were purchased from a research-grade porcine tissue vendor (Sierra Medical Inc.).

### Preparation of the barnacle-inspired paste

To prepare a bioadhesive, 30 w/w % acrylic acid, 2 w/w % chitosan, 1 w/w % AAc-NHS ester, and 0.5 w/w % α-ketoglutaric acid were dissolved in deionized water. The precursor solution was then filtered with 0.4 µm sterile syringe filters and poured on a glass mold with 500-µm spacers. The bioadhesive was cured in a UV chamber (284 nm, 10 W power) for 60 min and completely dried under nitrogen flow for 24 h. The dry bioadhesive was sealed in plastic bags with desiccants (silica gel packets) and stored in – 20 °C before use. To aid visualization of the barnacle-inspired paste for confocal microscope images, 0.2 w/w % fluorescein-labeled chitosan (FITC-chitosan) was further added into the precursor solution before curing. To prepare gelatin-based barnacle-inspired paste, 2 w/w % chitosan in the precursor solution was replaced with 10 w/w % gelatin.

To prepare bioadhesive microparticles, the dry bioadhesive was cut into small pieces and added into a container of a cryogenic grinder (CryoMill, Retsch), followed by cryogenic grinding process (30 Hz frequency for 2 min). The barnacle-inspired paste was prepared by thoroughly mixing the bioadhesive microparticles and a hydrophobic matrix. The prepared barnacle-inspired paste was sealed in plastic bags with desiccant (silica gel packets) and stored in – 20 °C before use. Unless otherwise specified, the silicone oil with viscosity of 5 cSt and the 1:1 mass ratio (equivalent to *ϕ* = 0.4) between the bioadhesive microparticles and the silicone oil were used.

### Mechanical tests

Unless otherwise indicated, all tissues were adhered by the barnacle-inspired paste after covering with blood or gastric juice followed by 5 s pressing (with 10 kPa pressure applied by either mechanical testing machine or equivalent weight). Unless otherwise indicated, all mechanical tests on adhesion samples were performed 6 h after initial pressing to ensure equilibrium swelling of the adhered barnacle-inspired paste. The application of commercially-available products followed the provided manual for each product.

For pull-off tests, porcine heart was cut with a surface area of 1 cm^2^ and thickness of 5 mm. On one side, the porcine heart was adhered to a glass container filled with DMEM or heparinized porcine blood bath by using a cyanoacrylate glue (Krazy Glue™). On another side, the porcine heart was adhered to an aluminum fixture by using the cyanoacrylate glue and the surface of the tissue was covered by the bioadhesive microparticles without or with varying matrix materials. Note that DMEM bath was used to mimic body fluids^38^. The adhesive-covered porcine heart was pressed against the tissue submerged in the bath at varying pressure by using a mechanical tester (2.5 kN load-cell, Zwick/Roell Z2.5) for 5 s. The adhered tissues were then pulled by lifting the aluminum fixture and the maximum tensile force was measured as the pull-off force.

To measure interfacial toughness, the adhered samples with 2.5 cm in width were prepared and tested by the standard 180-degree peel test (ASTM F2256) with the mechanical tester (2.5 kN load-cell, Zwick/Roell Z2.5). All tests were conducted with a constant peeling speed of 50 mm min^-1^. The measured force reached a plateau as the peeling process entered the steady-state. Interfacial toughness was determined by dividing two times of the plateau force (for 180-degree peel test) with the width of the tissue sample (**Supplementary Fig. 11a**). Poly(methyl methacrylate) (PMMA) film (50 µm thickness, Goodfellow) was applied by using the cyanoacrylate glue as a stiff backing for the tissues.

To measure shear strength, the adhered samples with an adhesion area of 2.5 cm in width and 1 cm in length were prepared and tested by the standard lap-shear test (ASTM F2255) with the mechanical tester. All tests were conducted with a constant tensile speed of 50 mm min^-1^. Shear strength was determined by dividing the maximum force by the adhesion area (**Supplementary Fig. 11b**). PMMA film was applied using the cyanoacrylate glue as a stiff backing for the tissues.

To measure tensile strength, the adhered samples with adhesion area of 2.5 cm in width and 2.5 cm in length were prepared and tested by the standard tensile test (ASTM F2258) with the mechanical tester (**Supplementary Fig. 11c**). All tests were conducted with a constant tensile speed of 50 mm min^-1^. Tensile strength was determined by dividing the maximum force with the adhesion area. Aluminum fixtures were applied by using the cyanoacrylate glue to provide grips for the tensile tests.

To measure burst pressure, a porcine aorta with area of 2.5 cm in width and 2.5 cm in length was prepared and tested by the standard burst pressure test (ASTM F2392-04) (**Fig. 5b**). A 3-mm hole was introduced to the porcine aorta by using a biopsy punch (Dynarex). The punctured porcine aorta was then covered with heparinized porcine blood and sealed by using various adhesives (the barnacle-inspired paste, suture (Ethicon 4-0 Vicryl), Surgiflo®, Tisseel, Tachosil®). 30 min after the sealing, pressure was applied to the sealed porcine aorta by pumping phosphate buffered saline (PBS) using a syringe pump at a flow rate of 2 ml min^-1^. The maximum pressure was recorded as the burst pressure by using a digital pressure gauge (Omega).

### Microscope imaging

Scanning electron microscope (SEM) images of the cryogenically ground bioadhesive microparticles were taken by using a SEM facility (JSM-6010LA, JEOL) with 5 nm gold sputtering to enhance image contrasts. Confocal microscope images of the barnacle-inspired paste were obtained by an upright confocal microscope (SP8, Leica) with 490 nm excitation wavelength for FITC.

### Contact angle measurement

The dry bioadhesive or a porcine skin tissue were bonded on a glass substrate and the contact angle of silicone oil and porcine blood was measured by using a contact angle apparatus (Ramé-Hart). The contact angle measurements were conducted at room temperature (23-26 °C) with relative humidity of 35 %. The interfacial energies between silicone oil/air, blood/air, and silicone oil/blood were obtained from the reported values in the literatures^39–41^.

### *Ex vivo* tests

All *ex vivo* experiments were reviewed and approved by the Committee on Animal Care at the Massachusetts Institute of Technology. For hemostatic sealing of a bleeding aorta, a porcine aorta was connected with a heparinized porcine blood bath and a pump via silicone tubes to generate closed-loop blood flow at 150 mmHg pressure (**Fig. 5a**). A 3-mm diameter hole was made to the porcine aorta by a biopsy punch (Dynarex). To form hemostatic seal, 1 ml of the barnacle-inspired paste was injected on a surgical gauze with the size of 2.5 cm in width and 2.5 cm in length to aid handling, and then gently pressed onto the bleeding site for 5 s. The sealed porcine aorta was kept for 6 h at room temperature with continuous blood flow to monitor the hemostatic sealing made by the barnacle-inspired paste. 0.01 w/v % sodium azide solution (in PBS) was sprayed on the porcine aorta to avoid tissue dehydration and degradation during the test. After the test, the blood bath was filtered by using a 100-µm mesh to check the presence of free-floating bioadhesive microparticles in the blood.

### *In vitro* biocompatibility evaluation

*In vitro* biocompatibility tests were conducted by using the barnacle-inspired paste-conditioned media for cell culture. To prepare the barnacle-inspired paste-conditioned or Coseal-conditioned media, 0.5 ml of the barnacle-inspired paste or Coseal were incubated in 10 mL DMEM supplemented with 10 v/v % fetal bovine serum (FBS) and 100 U ml^-1^ penicillin–streptomycin at 37 °C for 24 h. The supplemented DMEM without incubating the barnacle-inspired paste was used as a control. Rat embryonic cardiomyocytes (H9c2(2-1), ATCC) were plated in confocal dish (20-mm diameter) at a density of 0.5 × 10^5^ cells (*n* = 4 per each group). The cells were then treated with the barnacle-inspired paste-conditioned media and incubated at 37 °C for 24 h in 5 % CO_2_. The cell viability was determined by a LIVE/DEAD viability/cytotoxicity kit for mammalian cells (Thermo Fisher Scientific). A laser confocal microscope (SP 8, Leica) was used to image live cells with excitation/emission at 495nm/515nm, and dead cells at 495nm/635nm, respectively. The cell viability was calculated by counting the number of live (green fluorescence) and dead (red fluorescence) cells by using ImageJ (version 2.1.0).

### *In vivo* biocompatibility and biodegradability evaluation

All animal surgeries were reviewed and approved by the Committee on Animal Care at the Massachusetts Institute of Technology. Female Sprague Dawley rats (225-250 g, Charles River Laboratories) were used for all *in vivo* studies.

Prior to implantation, the barnacle-inspired paste was prepared using aseptic techniques and were further sterilized for 3 h under UV light. For implantation in the dorsal subcutaneous space, rats were anaesthetized using isoflurane (1–2% isoflurane in oxygen) in an anesthetizing chamber. Anesthesia was maintained using a nose cone. Back hair was removed and the animals were placed over a heating pad for the duration of the surgery. The subcutaneous space was accesed by a 1-2 cm skin incision per implant in the center of the animal’s back. To create space for implant placement, blunt disection was performed from the incision towards the animal shoulder blades. Either 0.5 ml of the barnacle-inspired paste (*n* = 4 for each endpoint) or a comparable volume of commercially-available tissue adhesive (Coseal, *n* = 4 for each endpoint) were placed in the subcutaneous pocket created above the incision. The incision was closed using interrupted sutures (4-0 Vicryl, Ethicon) and 3-6 ml of saline were injected subcutaneously. Up to four implants were placed per animal ensuring no overlap between each subcutaneous pocket created. After 1 day, 3 days, 1 week, or 2 weeks following the implantation, the animals were euthanized by CO_2_ inhalation. Subcutaneous regions of interest were excised and fixed in 10 % formalin for 24 h for histological and immunofluorescence analyses.

### *In vivo* hemostatic sealing of liver

For hemostatic sealing of the hepatic injury, the animals were anaesthetized using isoflurane (1–3% isoflurane in oxygen) in an anesthetizing chamber. Abdominal hair was removed and the animals were placed over a heating pad for the duration of the surgery. The liver was exposed via a laparotomy. A 5-mm diameter and 2-mm depth injury was made to the heart by using a biopsy punch (Dynarex). To form hemostatic sealing, 0.5 ml of the barnacle-inspired paste was injected onto the bleeding site and then gently pressed onto the punctured hole using a surgical spatula for 5 s (*n* = 4). For commercially-available products, Surgicel (Ethicon) with size of 20 mm in length and 20 mm in width (*n* = 4) or 2 ml of Coseal (Baxter) (*n* = 4) were used. For injury group, no hemostasis was performed (*n* = 4). The amount of blood loss until hemostasis and the time to hemostasis were recorded for each group. After the hemostatic sealing was confirmed, the incision was closed using interrupted sutures (4-0 Vicryl, Ethicon) and 3-6 ml of saline were injected subcutaneously. After 2 weeks following the implantation, blood was collected for the blood analysis and the animals were euthanized by CO_2_ inhalation. Livers with the implants were excised and fixed in 10 % formalin for 24 h for histological and immunofluorescence analyses.

### *In vivo* hemostatic sealing of heart

For hemostatic sealing of the full thickness ventricular injury, the animals were anaesthetized using isoflurane (1–3% isoflurane in oxygen) in an anesthetizing chamber. Chest hair was removed. Endotracheal intubation was performed, and the animals were connected to a mechanical ventilator (Model 683, Harvard Apparatus) and placed over a heating pad for the duration of the surgery. The heart was exposed via a thoracotomy and the pericardium was removed using fine forceps. For measurement of intraventricular blood pressure, a pressure-volume (PV) catheter (SPR-838, Millar) was inserted into the left ventricle (LV) via apical stick to monitor LV blood pressure during the test. A 2-mm diameter injury was made to the left or right ventricular wall of the heart by using a biopsy punch (Dynarex). To form hemostatic sealing, 0.25 ml of the barnacle-inspired paste was injected onto the bleeding site and then gently pressed onto the punctured hole using a surgical spatula for 5 s (*n* = 5). For commercially-available products, Surgicel (Ethicon) with size of 20 mm in length and 20 mm in width (*n* = 4) or 2 ml of Coseal (Baxter) (*n* = 4) were used. For injury group, no hemostasis was performed (*n* = 4). The amount of blood loss until hemostasis and the time to hemostasis were recorded up to 300 s for each group. After the hemostatic sealing was confirmed, the incision was closed using interrupted sutures (4-0 Vicryl, Ethicon) and 3-6 ml of saline were injected subcutaneously. For groups failed to form hemostasis until 300 s, the animals were euthanized by exsanguination. After 2 weeks following the implantation, blood was collected for the blood analysis and the animals were euthanized by CO_2_ inhalation. Hearts with the implants were excised and fixed in 10 % formalin for 24 h for histological and immunofluorescence analyses.

### Histological processing

Fixed tissue samples were placed into 70 % ethanol and submitted for histological processing and hematoxylin and eosin (H&E) or Masson’s trichrome staining at the Hope Babette Tang (1983) Histology Facility in the Koch Institute for Integrative Cancer Research at the Massachussets Institute of Technology. Histological assessment was performed by a blinded pathologist and representative images of each group were shown in the corresponding figures.

### Immunofluorescence analysis

The expression of targeted proteins (αSMA, Collagen I, CD68, CD3) were analyzed after the immunofluorescence staining of the collected tissues. Before the immunofluorescence analysis, the paraffin-imbedded fixed tissues were sliced and prepared into slides. The slides were deparaffinized and rehydrated to deionized water. Antigen retrieval was performed using steam method during which the slides were steamed in IHC-Tek Epitope Retrieval Solution (IW-1100) for 35 min and then cooled for 20 min. Then the slides were washed in three changes of PBS for 5 min per each cycle. After washing, the slides were incubated in primary antibodies (1:200 mouse anti-αSMA for fibroblast (ab7817, Abcam); 1:200 mouse anti-CD68 for macrophages (ab201340, Abcam); 1:100 rabbit anti-CD3 for T-cells (ab5690, Abcam); 1:200 rabbit anti-collagen-I for collagen (ab21286, Abcam)) diluted with IHC-Tek Antibody Diluent for 1 h at room temperature. The slides were then washed three times in PBS and incubated with Alexa Fluor 488 labeled anti-rabbit or anti-mouse secondary antibody (1:200, Jackson Immunoresearch) for 30 min. The slides were washed in PBS and then counterstained with propidium iodide solution for 20 min. A laser confocal microscope (SP 8, Leica) was used for image acquisition. ImageJ (version 2.1.0) was used to quantify the fluorescence intensity of expressed antibodies. All the images were transformed to the 8-bit binary images, and the fluorescence intensity was calculated with normalized analysis. All analyses were blinded with respect to the experimental conditions.

### Statistical analysis

MATLAB (version R2018b) was used to assess the statistical significance of all comparison studies in this work. Data distribution was assumed to be normal for all parametric tests, but not formally tested. In the statistical analysis for comparison between multiple samples, one-way ANOVA followed by Tukey’s multiple comparison test were conducted with the threshold of \**p* ≤ 0.05, \*\**p* ≤ 0.01, and \*\*\**p* ≤ 0.001. In the statistical analysis between two data groups, the two-sample Student’s *t*-test was used, and the significance threshold was placed at \**p* ≤ 0.05, \*\**p* ≤ 0.01, and \*\*\**p* ≤ 0.001.

## Data Availability

The main data supporting the findings in this study are available within the paper and its Supplementary Information. The raw and analyzed datasets are too numerous to be readily shared publicly. Source data for the figures are available for research purposes from the corresponding author on reasonable request.

## Code Availability

No custom code is used in this study.

## Acknowledgments

The authors thank the Koch Institute Swanson Biotechnology Center for technical support, specifically K. Cormier and the Histology Core for the histological processing, and Dr. R. Bronson at Harvard Medical School for the histological analyses. This work is supported by National Science Foundation (CMMI-1661627), National Institute of Health (1-R01-HL153857-01), and the U.S. Army Research Office through the Institute for Soldier Nanotechnologies at MIT (W911NF-13-D-0001). H.Y. acknowledges the financial support from Samsung Scholarship.

## Author Contributions

H.Y. conceived the idea and developed the materials and method for the barnacle-inspired paste. H.Y., X.M., and C.S.N. designed the *in vitro* and *ex vivo* experiments. H.Y., X.M., and J.W. conducted the *in vitro* and *ex vivo* experiments. H.Y., J.W., C.E.V., E.T.R., C.S.N. designed the *in vivo* experiments. J.W., H.Y., and C.E.V. conducted the *in vivo* experiments. H.Y., C.S.N., and X.Z. analyzed the results and wrote the manuscript with inputs from all authors.

