## Supplementary Information for "Barnacle-Inspired Paste for Instant Hemostatic Tissue Sealing"

**This PDF file includes:**

Supplementary Texts
Supplementary Figs. 1 to 24
Supplementary References
Captions for Supplementary Videos 1 to 5

**Other Supplementary Materials for this manuscript include the following:**

Supplementary Videos 1 to 5

### Stability of the Barnacle-Inspired Paste

To ensure stability of the barnacle-inspired paste, the bioadhesive microparticles should form a stable granular suspension with the silicone oil matrix. To satisfy this condition, it is essential that the bioadhesive microparticles should be preferentially wetted by the silicone oil rather than being phase-separated with the oil matrix.

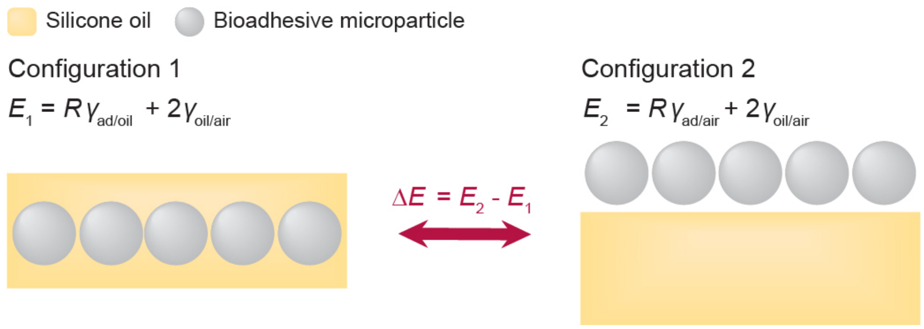


To evaluate this requirement, we compare the total surface energy of the individual wetting configurations (see above schematic illustration). Configuration 1 refers to the state where the bioadhesive microparticles are completely wetted by the silicone oil matrix, and thus provide the stable barnacle-inspired paste. Configuration 2 refers to the state where the bioadhesive microparticles are phase-separated with the silicone oil matrix, and thus result in the unstable barnacle-inspired paste. Here we consider the criterion whether the stable barnacle-inspired paste (Configuration 1) is energetically stable or not (i.e., lower total energy). To meet this criterion, the Configuration 1 should always have a lower energy state than the Configurations 2, that is,

$$\Delta E = E_2 - E_1 > 0 \tag{S1}$$

Eq. (S1) is equivalent to

$$R\gamma_{\mathrm{oil/air}} cos\theta_{\mathrm{oil/ad}} > 0 \tag{S2}$$

where $R$ is the roughness factor representing the ratio of the actual and projected surface areas of the bioadhesive microparticles, $\gamma_{\mathrm{A/B}}$ represents the interfacial energy between A and B, and $\theta_{\mathrm{A/B}}$ represents the contact angle of A on B (subscript "ad" represents bioadhesive microparticle). Note that we take $R$ as $\pi$ for the bioadhesive microparticles in the barnacle-inspired paste based on the first-order approximation of tightly placed spherical particles with the same diameter. By plugging the corresponding values in the Eq. (S2) ($R = \pi$, $\gamma_{\mathrm{oil/air}} = 20.9$ mN m$^{-1}$, $\theta_{\mathrm{oil/ad}} = 4.5°$) it is evident that inequality in the Eq. (S2) is satisfied. Hence, the bioadhesive microparticles and the silicone oil matrix can form the stable barnacle-inspired paste.

### Rheological Property of the Barnacle-Inspired Paste

As a volume fraction of the bioadhesive microparticles in the barnacle-inspired paste increases, the barnacle-inspired paste undergoes transition from a fluidic to a thixotropic state (Supplementary Fig. 8). This transition stems from a significant increase in viscosity of the barnacle-inspired paste with higher volume fraction of the bioadhesive microparticles, which can be understood based on the solid particle suspension theories[1-3]. As the volume fraction $\phi$ of the particles increases, the distance between the nearest neighboring particles decreases. Hence, an increase in $\phi$ leads to higher resistance for matrix fluid to flow through particles, resulting in higher viscosity $\eta_r$. As $\phi$ further increases, the distance between the particles continues to decrease until it reaches a maximum packing fraction $\phi_{\mathrm{m}}$, beyond which the particles are in jammed state and cannot flow anymore. As a result, $\eta_r$ of the particle suspension divergently increases as $\phi$ approaches $\phi_{\mathrm{m}}$ (Ref. 2). Under the first-order approximation of monodispersed spherical microparticles, a classical theory by Krieger and Dougherty provides a relationship between $\eta_r$ and $\phi$ as[4]

$$\eta_r = \left(1 - \frac{\phi}{\phi_{\mathrm{m}}}\right)^{-B\phi_{\mathrm{m}}} \tag{S3}$$

where $B$ is the Einstein coefficient and $\phi_{\mathrm{m}}$ is the maximum packing fraction.

Experimental observations show that the barnacle-inspired paste exhibits gradual transition from a low viscosity fluid to a high viscosity thixotropic paste as the mass ratio between the bioadhesive microparticles and the silicone oil matrix increase from 1:3 to 1:0.5, whereas the barnacle-inspired paste starts to become a non-flowable solid when ratio of the bioadhesive microparticles to the silicone oil matrix becomes higher than 1:0.5 (Supplementary Fig. 8). With the given density of each component of the barnacle-inspired paste ($\rho_{\mathrm{particle}} = 1.39$ g mL$^{-1}$ and $\rho_{\mathrm{oil}} = 0.96$ g mL$^{-1}$), the transition mass ratio of 1:0.5 (bioadhesive microparticles:silicone oil) corresponds to $\phi = 0.58$, which is close to $\phi_{\mathrm{m}}$ of an homogeneously sheared assembly of spherical microparticles ($\phi_{\mathrm{m}} \sim 0.6$) (ref. 5) agreeing with the Eq. (S3).

### Compaction of the Barnacle-Inspired Paste

During hemostasis sealing, the bioadhesive microparticles in the barnacle-inspired paste are densely compacted into a non-flowable jammed layer (i.e., $\phi \rightarrow \phi_{\mathrm{m}}$) while the displaced silicone oil matrix from the compacted barnacle-inspired paste repels blood on the bleeding tissue. Since the barnacle-inspired paste takes the form of a granular suspension consisting of the bioadhesive microparticles and the silicone oil matrix, the required pressure to compact the barnacle-inspired paste is dependent to the viscosity of the silicone oil matrix. Under the first-order approximation of monodispersed spherical microparticles, a granular suspension theory by Guazzelli and Pouliquen provides a relationship for the required pressure to compact a granular suspension as[6]

$$P_{\mathrm{press}} \propto \frac{h}{d^2}(\phi_{\mathrm{m}} - \phi)U_x\eta_{\mathrm{m}} \tag{S4}$$

where $P_{\mathrm{press}}$ is the applied pressure, $h$ is the height of the suspension layer, $d$ is the particle diameter, $\phi_{\mathrm{m}}$ is the maximum packing fraction of the particles, $\phi$ is the particle volume fraction in the suspension, $U_x$ is the speed of the compaction, and $\eta_{\mathrm{m}}$ is the viscosity of the fluid matrix. Therefore, for the given barnacle-inspired paste ($d$, $h$, $\phi_{\mathrm{m}}$, and $\phi$) and $U_x$, more viscous silicone oil matrix requires the higher applied pressure to compact the barnacle-inspired paste to form hemostatic tissue sealing agreeing with the experimental data in Fig. 2f.

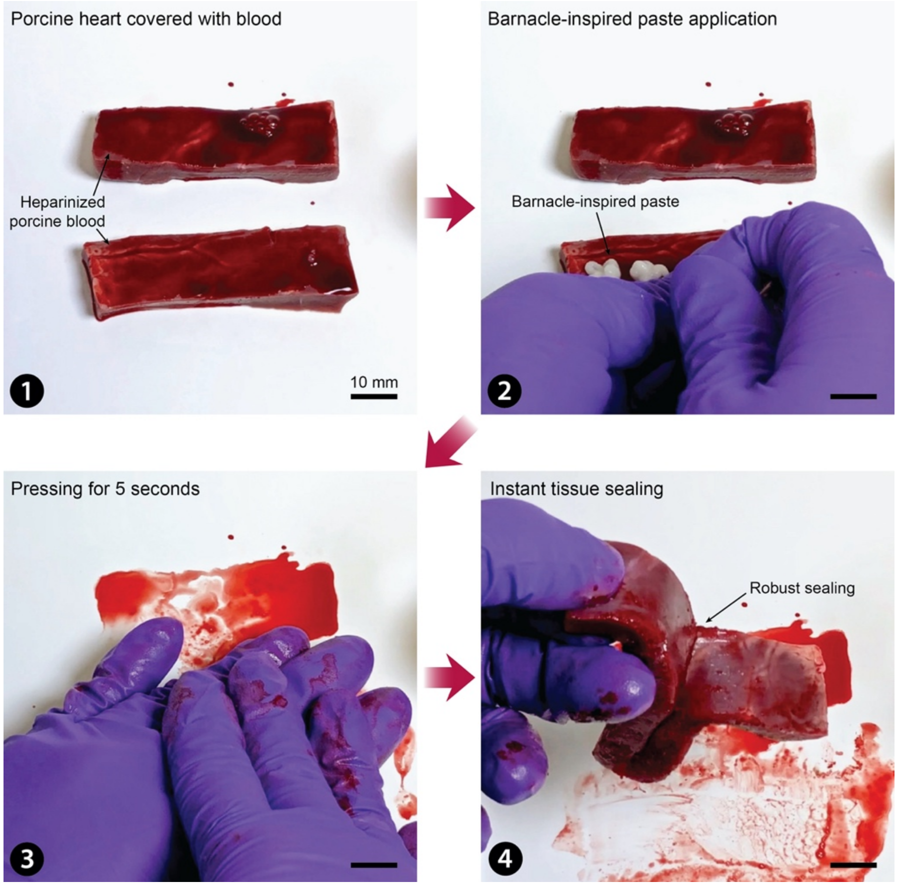


**Supplementary Fig. 1 | Instant hemostatic tissue sealing by the barnacle-inspired paste.** (1) Porcine heart covered with heparinized porcine blood; (2) the barnacle-inspired paste application; (3) gentle pressure application for 5 s; (4) instant hemostatic tissue sealing formation.

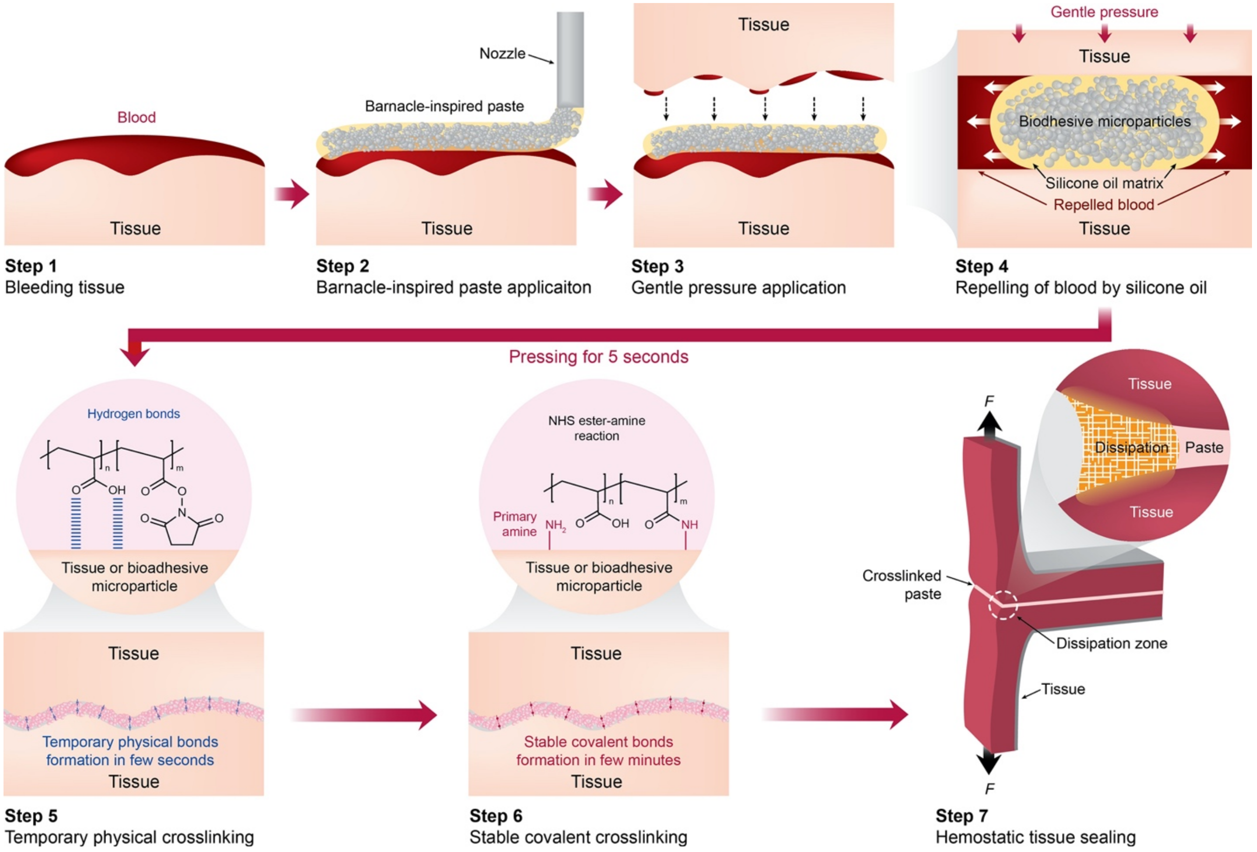


**Supplementary Fig. 2 | Schematic illustration for overall process of instant hemostatic tissue sealing by the barnacle-inspired paste.** (Step 1 and 2) The barnacle-inspired paste can be applied directly on bleeding injury without any other preparation process; (Step 3 and 4) Upon application of gentle pressure, the silicone oil matrix in the barnacle-inspired paste repels and clean blood from the bleeding injury; (Step 5 and 6) Simultaneously, the carboxylic acid groups in the bioadhesive microparticles form temporary physical crosslinks by hydrogen bonds, followed by the covalent crosslinking between the NHS ester groups and the primary amine groups with themselves and the tissue surfaces; (Step 7) the swollen and crosslinked barnacle-inspired paste provides robust hemostatic tissue sealing.

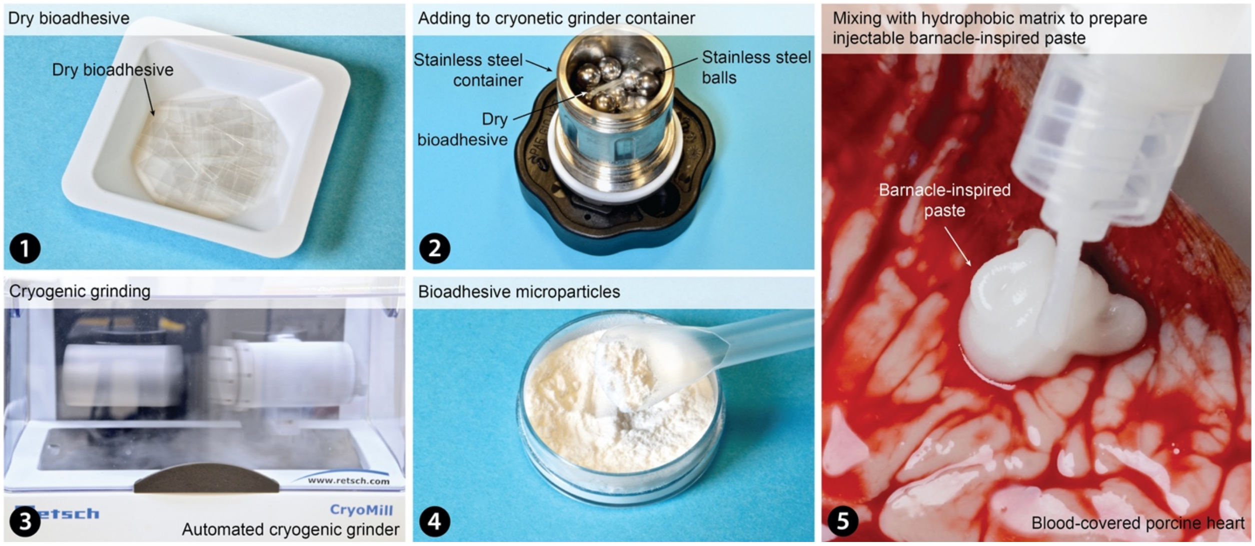


**Supplementary Fig. 3 | Preparation of the barnacle-inspired paste.** (1) Dry bioadhesive cut into small pieces; (2) Dry bioadhesive in stainless steel grinder container and balls; (3) Cryogenic grinding of dry bioadhesive; (4) Bioadhesive microparticle after the cryogenic grinding; (5) the barnacle-inspired paste after mixing the bioadhesive microparticle with the silicone oil matrix.

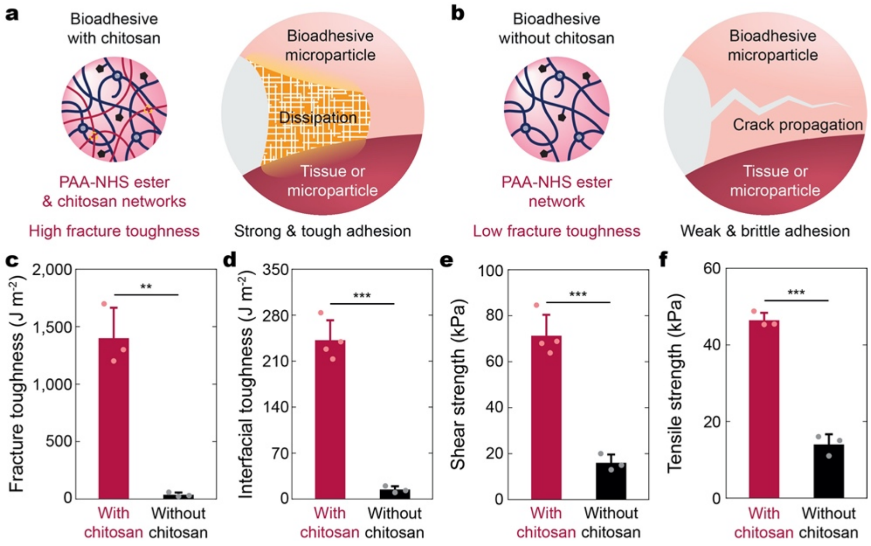


**Supplementary Fig. 4 | Effect of chitosan on the barnacle-inspired paste. a**,**b**, Schematic illustrations for the barnacle-inspired paste with chitosan (a) and without chitosan (b). **c-f**, Fracture toughness of the bioadhesive (c), interfacial toughness (d), shear strength (e), and tensile strength (f) of blood-covered porcine skin sealed by the barnacle-inspired paste with or without chitosan. Values in **c-f** represent the mean and the standard deviation ($n$ = 3-4 independent samples). Statistical significance and $p$ values are determined by two-sided Student $t$-test; ** $p \leq 0.01$; *** $p \leq 0.001$.

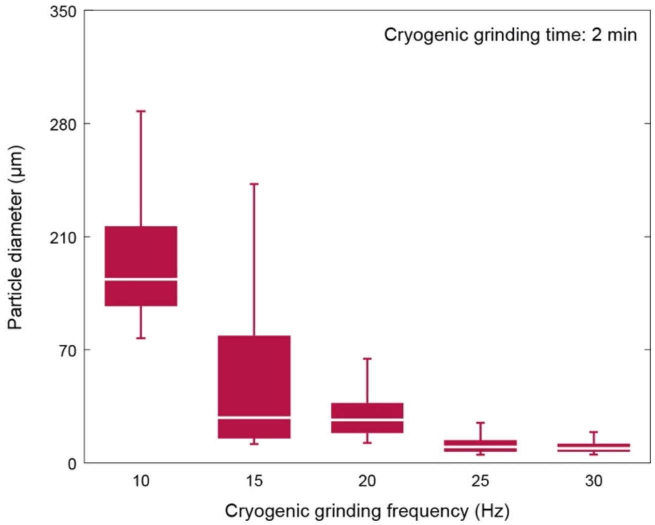


**Supplementary Fig. 5 | Size of bioadhesive microparticles under varying cryogenic grinding condition.** Cryogenic grinding frequency vs. bioadhesive microparticle diameter in a box plot. The whiskers correspond to the upper extreme and the lower extreme. The lines indicate the median and the error bars indicate the upper quartile and the lower quartile ($n$ = 50).

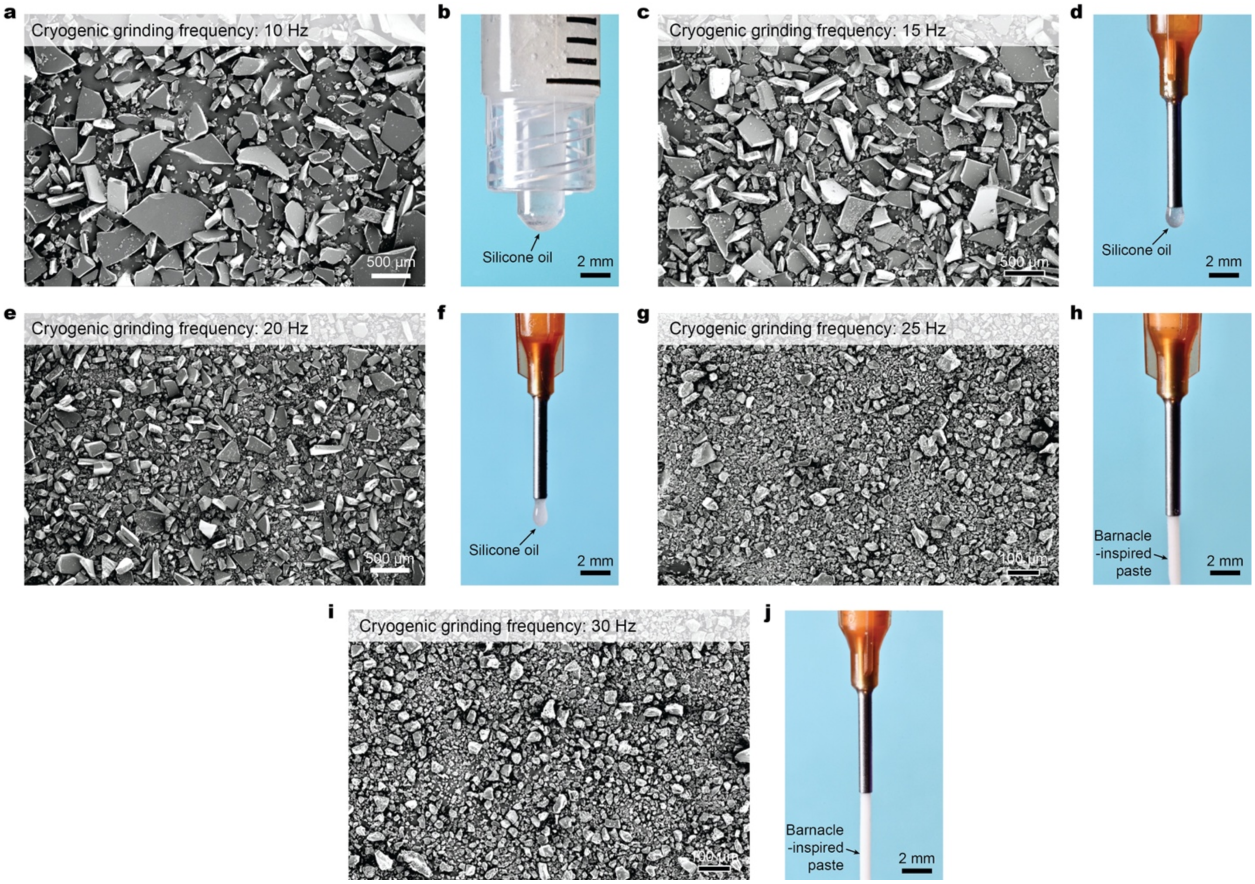


**Supplementary Fig. 6 | Injectability of the barnacle-inspired paste.** **a**,**b**, SEM image of the bioadhesive microparticles at 10 Hz cryogenic grinding frequency (a) and the corresponding barnacle-inspired paste injected through a syringe with 2.5-mm diameter (b). **c**,**d**, SEM image of the bioadhesive microparticles at 15 Hz cryogenic grinding frequency (c) and the corresponding barnacle-inspired paste injected through a nozzle with 1.2-mm diameter (d). **e**,**f**, SEM image of the bioadhesive microparticles at 20 Hz cryogenic grinding frequency (e) and the corresponding barnacle-inspired paste injected through a nozzle with 1.2-mm diameter (f). **g**,**h**, SEM image of the bioadhesive microparticles at 25 Hz cryogenic grinding frequency (g) and the corresponding barnacle-inspired paste injected through a nozzle with 1.2-mm diameter (h). **i**,**j**, SEM image of the bioadhesive microparticles at 30 Hz cryogenic grinding frequency (i) and the corresponding barnacle-inspired paste injected through a nozzle with 1.2-mm diameter (j). 2 min cryogenic grinding time is used for all cases.

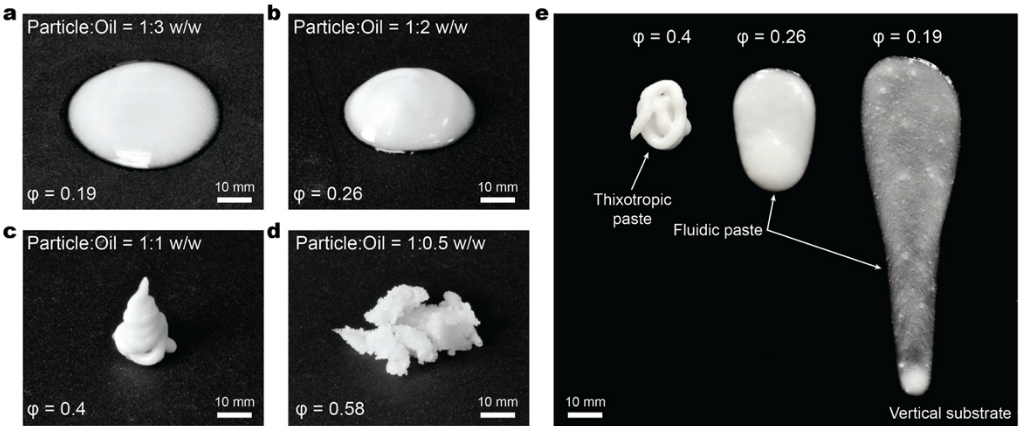


**Supplementary Fig. 7 | Rheological transition of the barnacle-inspired paste. a-d**, Images of the barnacle-inspired paste with the mass ratio between the bioadhesive microparticles and the silicone oil matrix at 1:3 (a), 1:2 (b), 1:1 (c), and 1:0.5 (d). **e**, Image of the barnacle-inspired paste with varying mixing ratio injected on a vertical substrate to visualize rheological property.

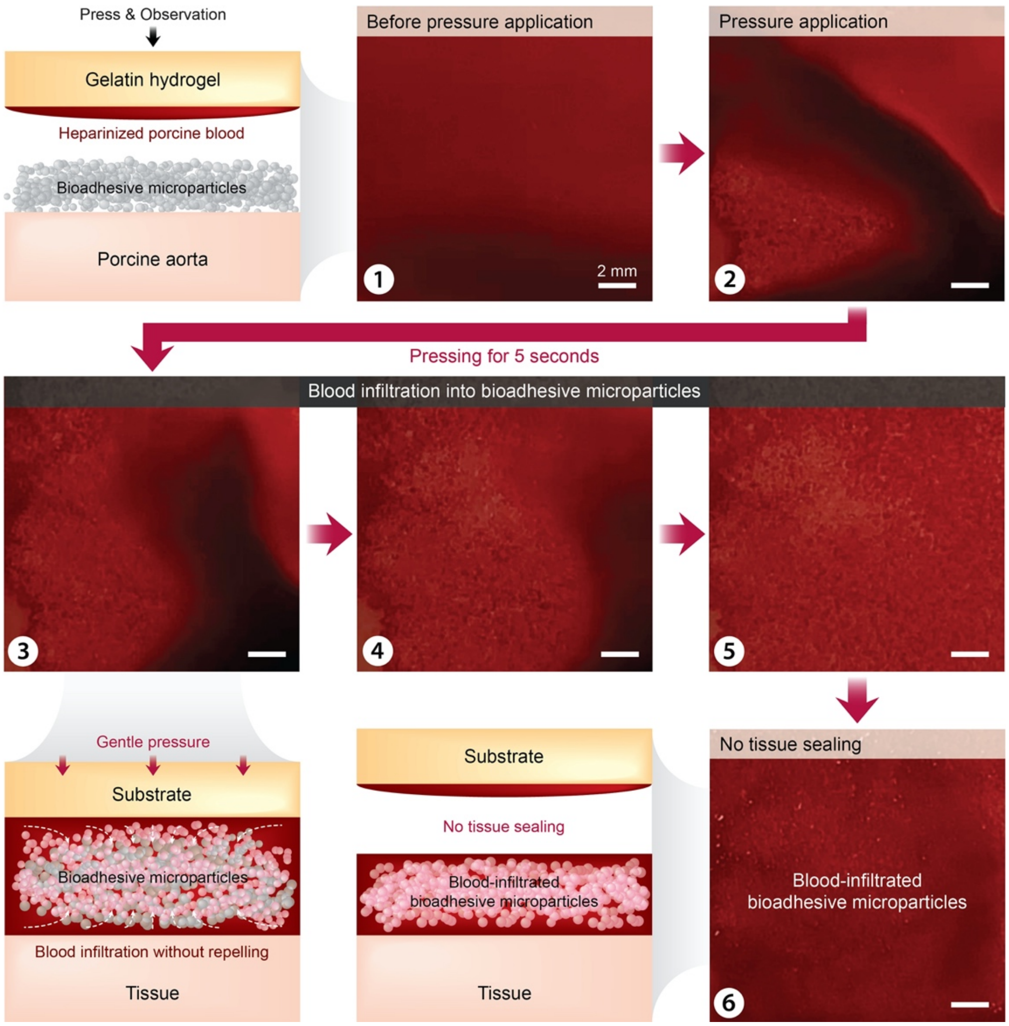


**Supplementary Fig. 8 | Blood incompatibility of the bioadhesive microparticles without hydrophobic silicone oil matrix.** (1) A heparinized porcine blood-covered gelatin hydrogel placed on top of a porcine aorta covered with a layer of bioadhesive microparticles without matrix; (2) The blood-covered gelatin hydrogel pressed on the tissue; (3) blood infiltrating into the bioadhesive microparticles; (4-5) swelling of the bioadhesive microparticles by the infiltrated blood; (6) no hemostatic sealing formation.

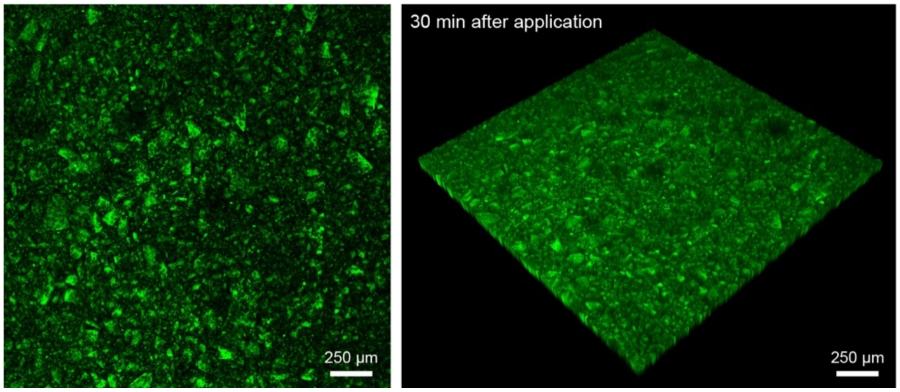


**Supplementary Fig. 9 | Densely compacted barnacle-inspired paste.** Confocal microscope images of the densely compacted barnacle-inspired paste between gelatin hydrogels after gentle pressure application for 5 s.

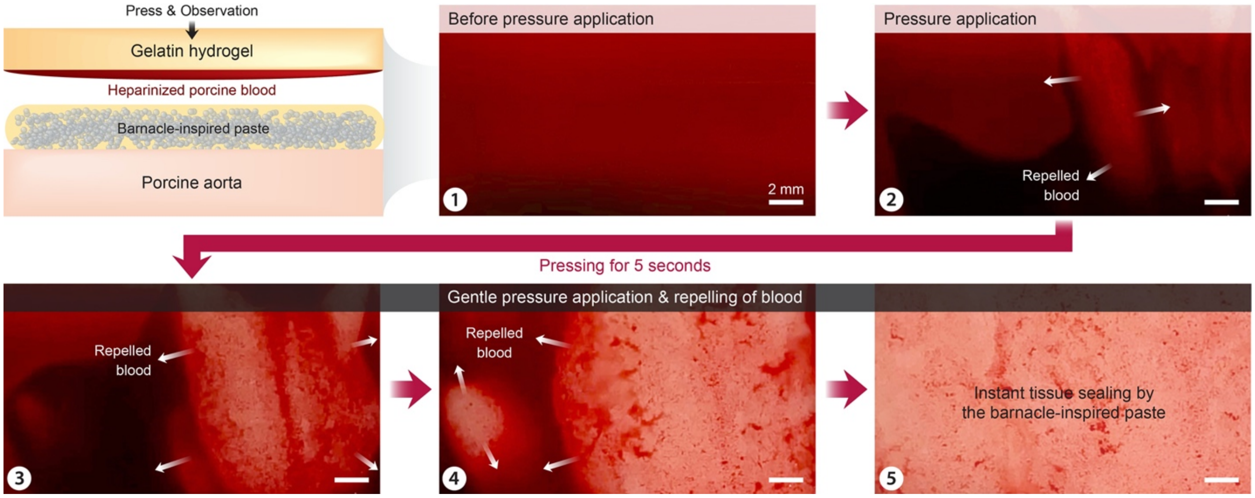


**Supplementary Fig. 10 | Blood repelling and hemostatic sealing by the barnacle-inspired paste.** (1) A heparinized porcine blood-covered gelatin hydrogel placed on top of a porcine aorta covered with a layer of barnacle-inspired paste; (2) The blood-covered gelatin hydrogel pressed on the tissue; (3) blood repelling by the barnacle-inspired paste; (4) cleaned tissue surface by the barnacle-inspired paste; (5) instant hemostatic sealing by the barnacle-inspired paste.

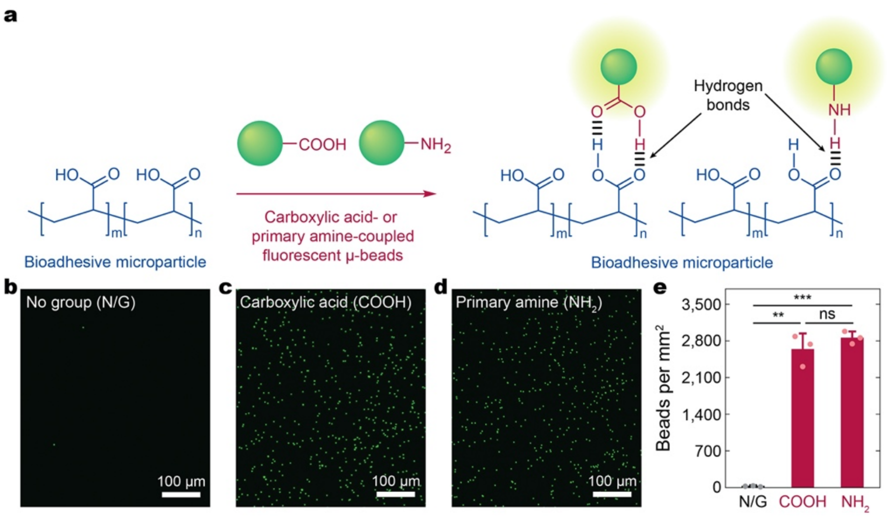


**Supplementary Fig. 11 | Hydrogen bonds-based physical crosslinking of the barnacle-inspired paste. a**, Schematic illustrations for hydrogen bonds formation between the barnacle-inspired paste and carboxylic acid (COOH) or primary amine ($NH_2$) coupled fluorescent microbeads. **b-d**, Images of the bioadhesive incubated in DMEM with pristine (b), carboxylic acid-coupled (c), and primary amine-coupled (d) fluorescent microbeads for 5 s. 3 independent experiments were conducted with similar results. N/G, no group. **e**, Number of pristine, carboxylic acid-coupled, and primary amine-coupled fluorescent microbeads on the bioadhesive per $mm^2$. Values in **e** represent the mean and the standard deviation ($n = 3$ independent samples). Statistical significance and $p$ values are determined by two-sided Student $t$-test; ns, not significant; ** $p \leq 0.01$; *** $p \leq 0.001$.

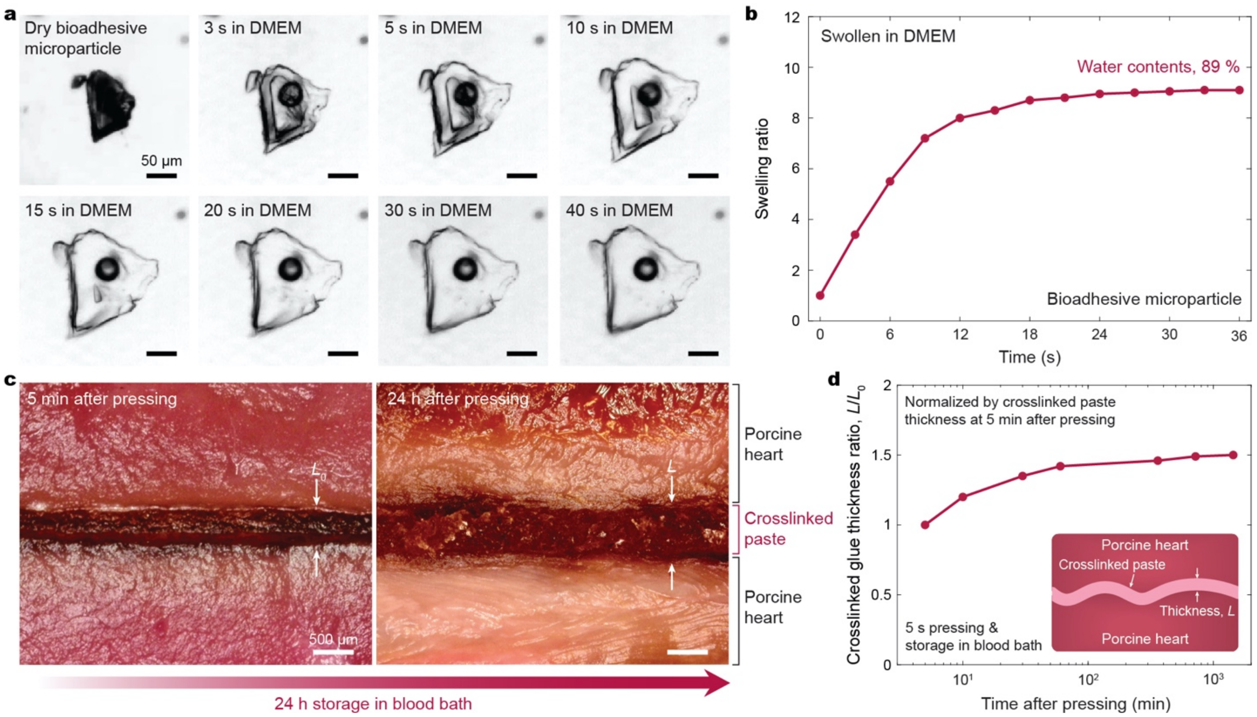


**Supplementary Fig. 12 | Swelling of the barnacle-inspired paste. a**,**b**, Images for swelling of a bioadhesive microparticle in DMEM (a) and the corresponding time vs. swelling ratio (b). **c**, Images of the cross-sectional view of blood-covered porcine heart sealed by the barnacle-inspired paste 5 min and 24 h after hemostatic sealing. $\boldsymbol{L_0}$, thickness of the barnacle-inspired paste 5 min after hemostatic sealing. **d**, Normalized thickness of the crosslinked barnacle-inspired paste between porcine heart over time. *L*, thickness of the barnacle-inspired paste at the current time.

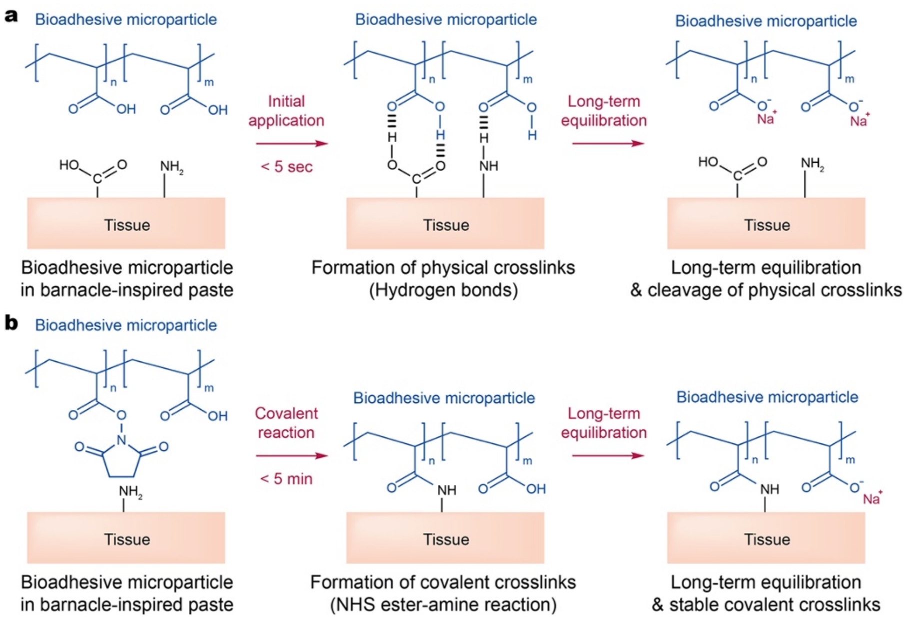


**Supplementary Fig. 13 | Adhesion mechanisms of the barnacle-inspired paste. a**, Schematic illustrations for the physical crosslinking-based adhesion between the barnacle-inspired paste and the tissue surface. **b**, Schematic illustrations for the covalent crosslinking-based adhesion between the barnacle-inspired paste and the tissue surface.

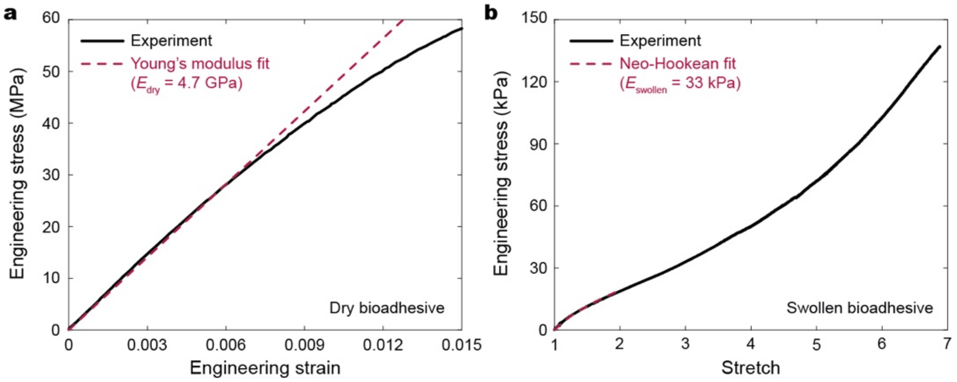


**Supplementary Fig. 14 | Mechanical property of bioadhesive in the barnacle-inspired paste.** **a**, Tensile engineering strain vs. engineering stress of the dry bioadhesive. $\boldsymbol{E}_{\mathbf{dry}}$, Young's modulus of the dry bioadhesive. **b**, Tensile stretch vs. engineering stress curve of the fully swollen bioadhesive in DMEM. $\boldsymbol{E}_{\mathbf{swollen}}$, Young's modulus of the swollen bioadhesive.

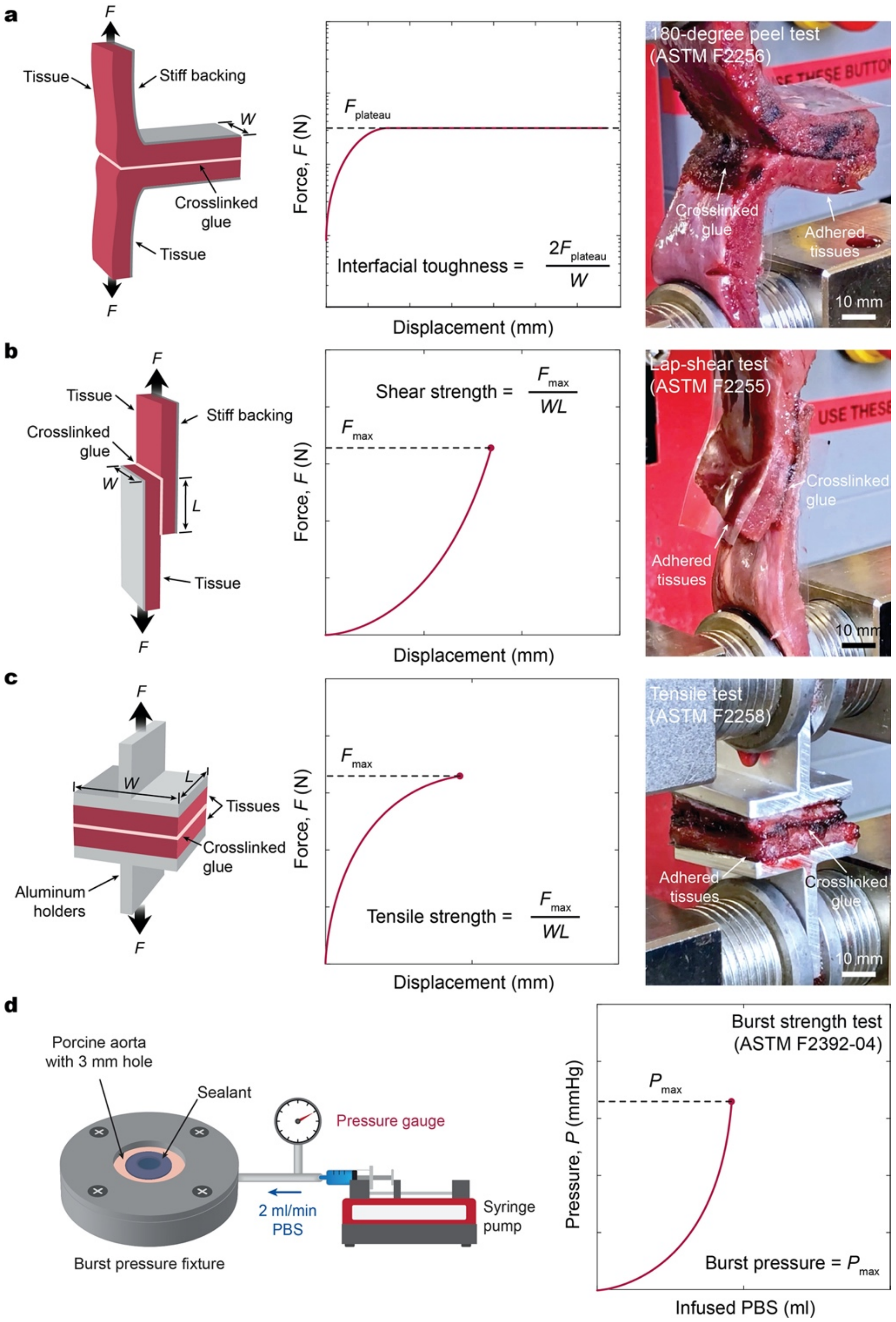


**Supplementary Fig. 15 | Mechanical test setups.** **a**, Test setup for interfacial toughness measurements based on the standard 180-degree peel test (ASTM F2256). **b**, Test setup for shear strength measurements based on the standard lap-shear test (ASTM F2255). **c**, Test setup for tensile strength measurements based on the standard tensile test (ASTM F2258). **d**, Test setup for burst pressure measurements based on the standard burst strength test (ASTM F2392-04).

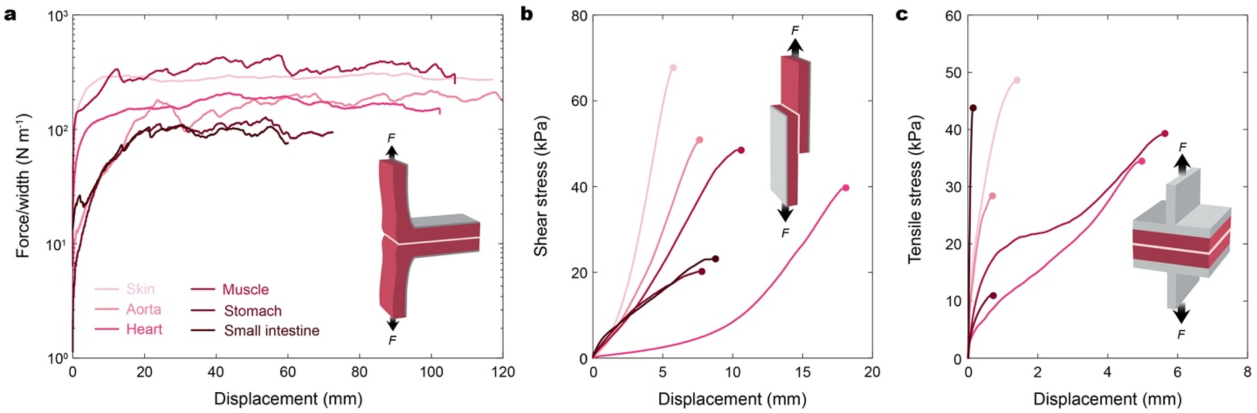


**Supplementary Fig. 16 | Representative curves for mechanical tests of various tissues sealed by the barnacle-inspired paste. a**, Force/width vs. displacement curves for the 180-degree peel tests of various tissues covered by a heparinized porcine blood or mucus sealed by the barnacle-inspired paste. **b**, Shear stress vs. displacement curves for the lap-shear tests of various tissues covered by a heparinized porcine blood or mucus sealed by the barnacle-inspired paste. **c**, Tensile stress vs. displacement curves for the tensile tests of various tissues covered by a heparinized porcine blood or mucus sealed by the barnacle-inspired paste.

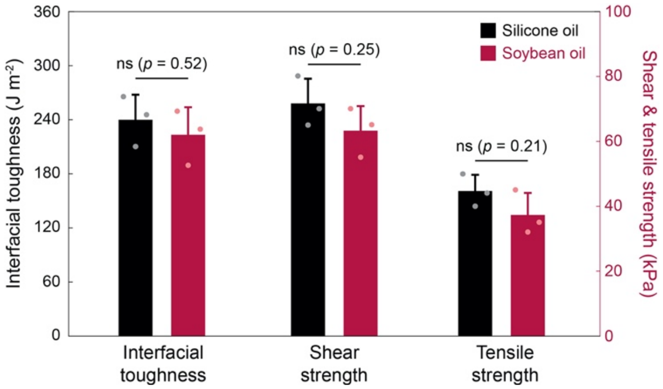


**Supplementary Fig. 17 | Adhesion performance of the barnacle-inspired paste based on various hydrophobic oil matrix.** Interfacial toughness, shear strength, and tensile strength between blood-covered porcine skin sealed by the barnacle-inspired paste based on silicone oil or soybean oil. $\boldsymbol{\phi}$ = 0.4 is used for both silicone oil- and soybean oil-based barnacle-inspired paste. Values in represent the mean and the standard deviation ($n$ = 3). Statistical significance and $p$ values are determined by two-sided Student $t$-test; ns, not significant.

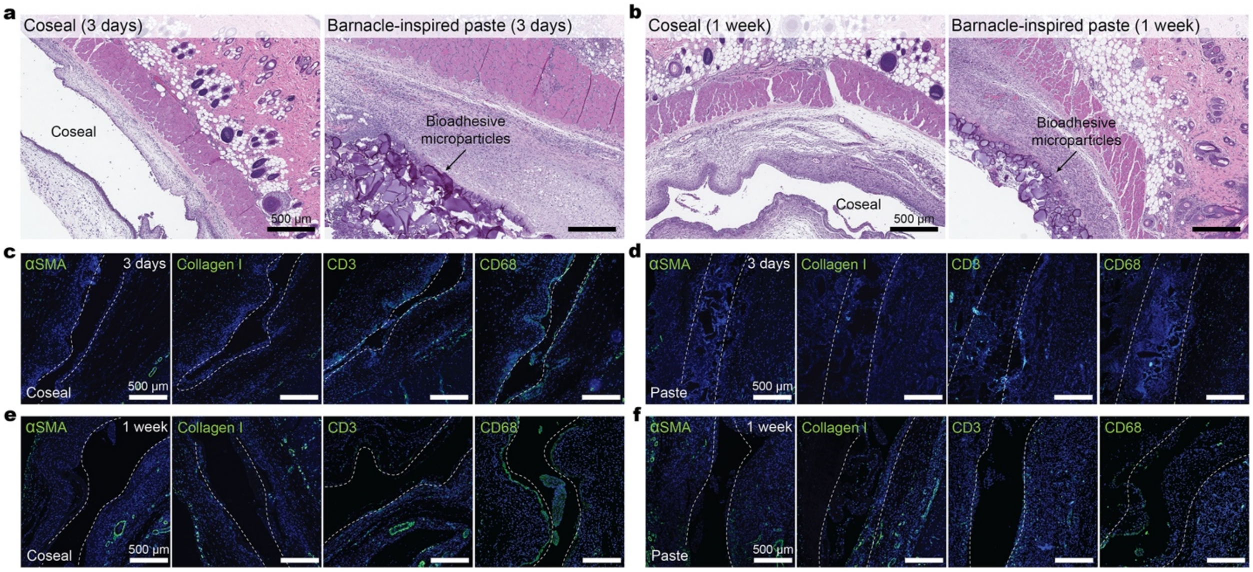


**Supplementary Fig. 18 | Histology and immunofluorescence images of the subcutaneous implants. a**,**b**, Representative histology images stained with hematoxylin and eosin (H&E) for Coseal and the barnacle-inspired paste after rat subcutaneous implantation for 3 days (a) and 1 week (b). 4 independent experiments were conducted with similar results. **c-f**, Representative immunofluorescence images of Coseal (c) and the barnacle-inspired paste (d) after rat subcutaneous implantation for 3 days; Coseal (e) and the barnacle-inspired paste (f) after rat subcutaneous implantation for 1 week. Cell nuclei are stained with 4′,6-diamidino-2-phenylindole (DAPI, Blue). Green fluorescence corresponds to the expression of fibroblast (αSMA), type 1 collagen (Collagen-I), T-cell (CD3), and macrophage (CD68), respectively.

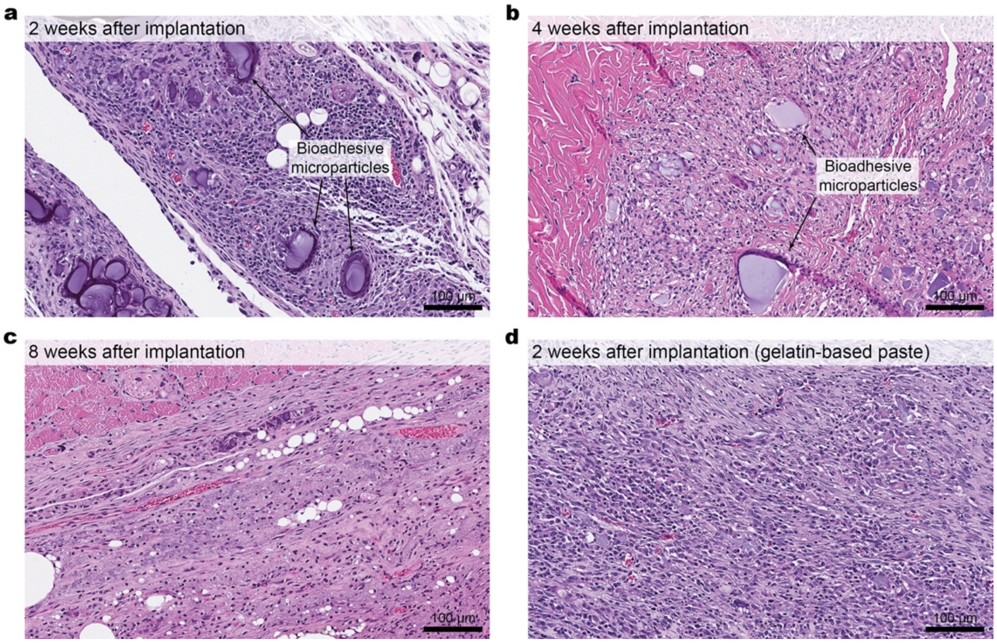


**Supplementary Fig. 19 | *In vivo* degradation of the barnacle-inspired paste. a-c**, Representative histology images stained with hematoxylin and eosin (H&E) for the barnacle-inspired paste after rat subcutaneous implantation for 2 weeks (a), 4 weeks (b), and 8 weeks (c). 4 independent experiments were conducted with similar results. **d**, Representative histology image stained with hematoxylin and eosin (H&E) for the gelatin-based barnacle-inspired paste after rat subcutaneous implantation for 2 weeks. 4 independent experiments were conducted with similar results.

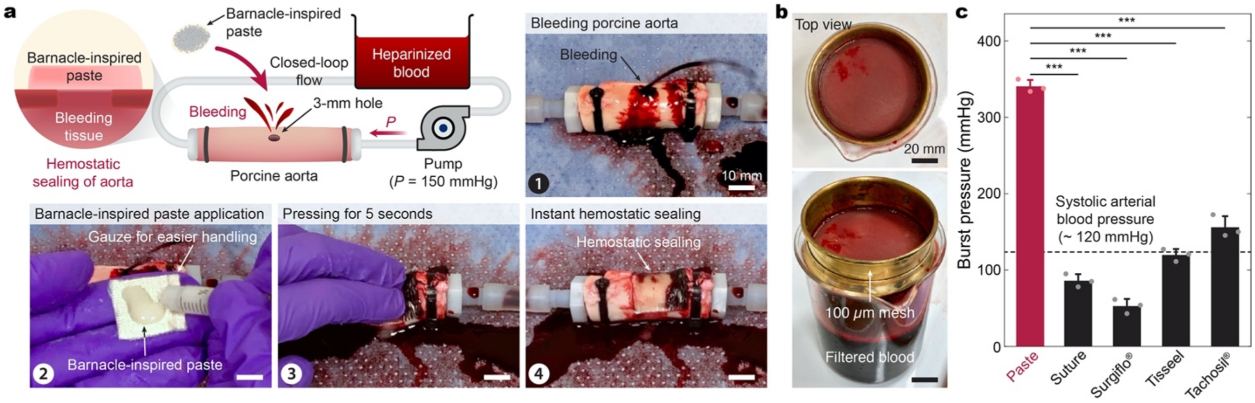


**Supplementary Fig. 20 | Instant and coagulation-independent hemostatic sealing of aorta by the barnacle-inspired paste. a**, Instant hemostatic sealing of an *ex vivo* porcine aorta by the barnacle-inspired paste. A heparinized porcine blood is used to ensure coagulation-independent hemostatic sealing. **b**, Images of a filtered porcine blood bath with a 100-µm mesh after 6 h continuous flow through a sealed *ex vivo* porcine aorta. **c**, Burst pressure of porcine aorta sealed by the barnacle-inspired paste and commercially-available products. Values in **c** represent the mean and the standard deviation ($n = 3$ independent samples). Statistical significance and $p$ values are determined by two-sided Student $t$-test; *** $p \leq 0.001$.

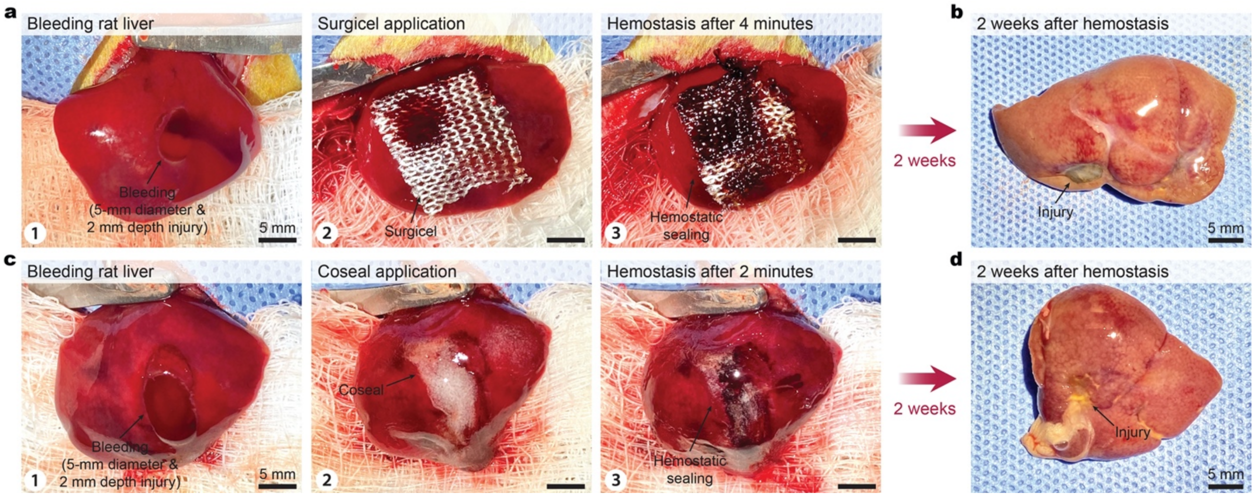


**Supplementary Fig. 21 | Hemostatic sealing of liver by commercially-available products. a**, Hemostatic sealing of a bleeding rat liver *in vivo* by Surgicel. **b**, Excised rat liver 2 weeks after hemostatic sealing by Surgicel. **c**, Hemostatic sealing of a bleeding rat liver *in vivo* by Coseal. **d**, Excised rat liver 2 weeks after hemostatic sealing by Coseal.

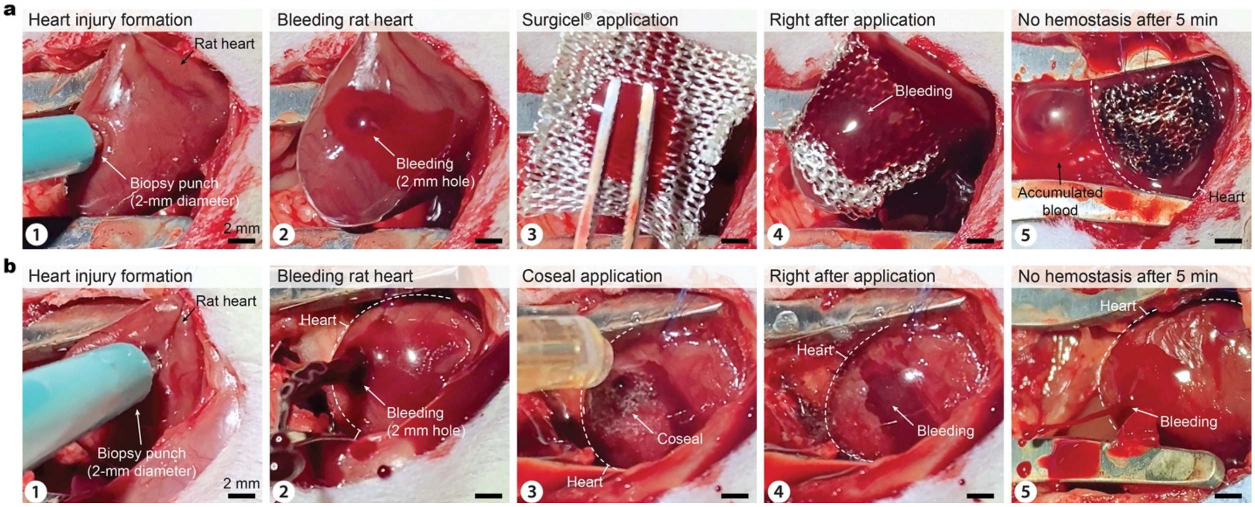


**Supplementary Fig. 22 | Hemostatic sealing of heart by commercially-available products. a**, Hemostatic sealing of a bleeding rat heart *in vivo* by Surgicel. **b**, Hemostatic sealing of a bleeding rat heart *in vivo* by Coseal.

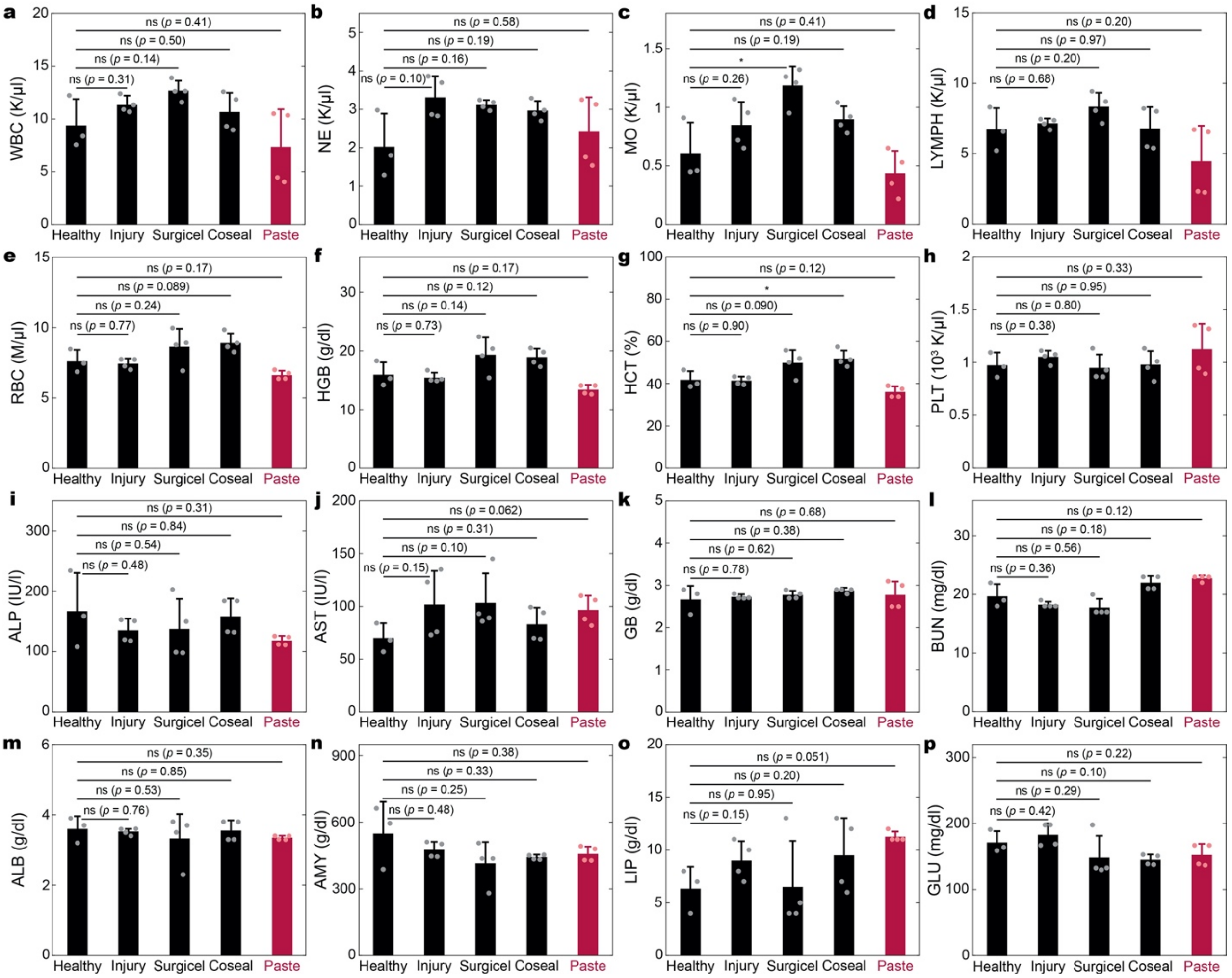

**Supplementary Fig. 23 | Blood analysis of the animals with hemostatic sealing of liver. a-h**, Complete blood count (CBC) of the healthy animals and the animals 2 weeks after hemostatic sealing of liver for white blood cell (WBC, a), neutrophil (NE, b), monocyte (MO, c), lymphocyte (LYMPH, d), red blood cell (RBC, e), hemoglobin (HGB, f), hematocrit (HCT, g), and platelet (PLT, h). **i-p**, Blood chemistry of the healthy animals and the animals 2 weeks after hemostatic sealing of liver for alkaline phosphatase (ALP, i), aspartate transaminase (AST, j), globulin (GB, k), blood urea nitrogen (BUN, l), albumin (ALB, m), amylase (AMY, n), lipase (LIP, o), and glucose (GLU, p). Values represent the mean and the standard deviation ($n$ = 4 independent samples). Statistical significance and $p$ values are determined by two-sided Student $t$-test; ns, not significant; * $p \leq 0.05$.

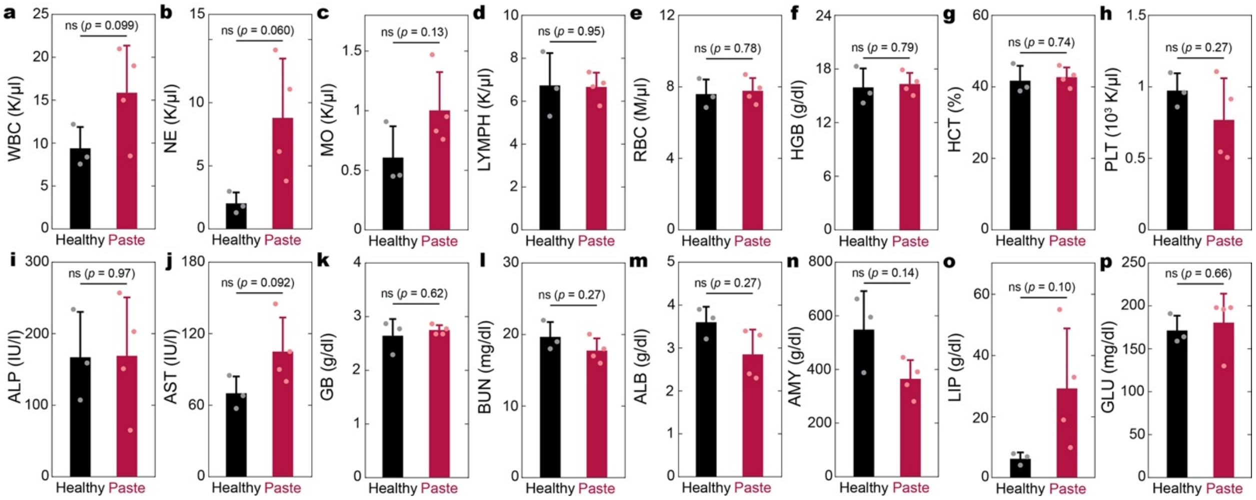


**Supplementary Fig. 24 | Blood analysis of the animals with hemostatic sealing of heart. a-h**, Complete blood count (CBC) of the healthy animals and the animals 2 weeks after hemostatic sealing of heart for white blood cell (WBC, a), neutrophil (NE, b), monocyte (MO, c), lymphocyte (LYMPH, d), red blood cell (RBC, e), hemoglobin (HGB, f), hematocrit (HCT, g), and platelet (PLT, h). **i-p**, Blood chemistry of the healthy animals and the animals 2 weeks after hemostatic sealing of heart for alkaline phosphatase (ALP, i), aspartate transaminase (AST, j), globulin (GB, k), blood urea nitrogen (BUN, l), albumin (ALB, m), amylase (AMY, n), lipase (LIP, o), and glucose (GLU, p). Values represent the mean and the standard deviation ($n$ = 4 independent samples). Statistical significance and $p$ values are determined by two-sided Student $t$-test; ns, not significant.

**Captions for Supplementary Videos**

**Supplementary Video 1 |** Overall process of adhesion formation between blood-covered tissues by the barnacle-inspired paste.

**Supplementary Video 2 |** Repelling of blood by the barnacle-inspired paste applied on a porcine aorta.

**Supplementary Video 3 |** Instant coagulation-independent hemostatic tissue sealing by the barnacle-inspired paste.

**Supplementary Video 4 |** Application of the commercially-available tissue adhesives to a bleeding *in vivo* rat heart.

**Supplementary Video 5 |** Instant hemostatic tissue sealing of a bleeding *in vivo* rat heart by the barnacle-inspired paste.
